# The Impact of The Hydroxymethyl-Cytosine Epigenetic Signature on DNA Structure And Function

**DOI:** 10.1101/2020.09.17.285452

**Authors:** Federica Battistini, Pablo D. Dans, Montserrat Terrazas, Chiara L. Castellazzi, Guillem Portella, Mireia Labrador, Núria Villegas, Isabelle Brun-Heath, Carlos González, Modesto Orozco

## Abstract

We present a comprehensive, experimental and theoretical study of the impact of 5-hydroxymethylation of DNA cytosine. Using molecular dynamics, biophysical experiments and NMR spectroscopy, we found that Ten-Eleven translocation (TET) dioxygenases generate an epigenetic variant with structural and physical properties not too different to those of 5-methylcytosine. Experiments and simulations demonstrate that 5-methyl-cytosine (mC) and 5-hydroxymethyl-cytosine (hmC) generally lead to more rigid duplexes with poorer circularization efficiencies and lower ability to form nucleosomes. In particular, we can rule out the hypothesis that hydroxymethylation reverts to unmodified cytosine physical properties, as hmC is even more rigid than mC. Thus, we do not expect dramatic changes in the chromatin structure induced by differences in physical properties between d(mCpG) and d(hmCpG). On the contrary, our simulations suggest that methylated-DNA binding domains (MBD), associated with repression activities, are very sensitive to the substitution d(mCpG)→ d(hmCpG), while MBD3 which has a dual activation/repression activity is not sensitive to the d(mCpG) → d(hmCpG) change. Overall, while changes in gene activity due to cytosine methylation are the result of the combination of stiffness-related chromatin reorganization and MBD binding, those associated to 5-hydroxylation of methylcytosine could be explained by a change in the balance of repression/activation pathways related to differential MBD binding.

## INTRODUCTION

Genomic expression is a finely controlled process regulated by a myriad of external and internal effectors which ensure that genetic information is expressed whenever and wherever required in the interest of the entire organism (1). External effectors are proteins or RNAs, which can regulate directly DNA transcription activation or repression. Internal effectors are chromatin chemical modifications (epigenetic changes) (2) that can affect DNA expression by recruiting activator or repressor proteins; or by controlling the accessibility of the transcriptional machinery to the target DNA sequences.

In developed organisms, such as mammals, DNA epigenetic changes are highly prevalent and most CpG steps (except for those in CpG islands) are methylated at the 5- position of the pyrimidine ring (3, 4). The presence of 5-methylcytosine (mC) has been traditionally related to gene inactivation,(5–8) but analysis of methylation changes in Leukaemia strongly suggests that the real connection between methylation and gene expression is probably more complex and differs in each individual gene (9). It is known that methylated DNA (met-DNA) recruits proteins containing methylated-DNA binding domains (MBD;10) triggering protein-specific responses. Furthermore, it has been described that DNA methylation modulates DNA accessibility, (11) mainly by making the DNA stiffer and less prone to wrap around nucleosomes (12, 13), leading to nucleosome shifts and accordingly to changes in the pattern of accessible and hidden sequences.

The portfolio of DNA epigenetic modifications in mammals has been recently expanded with 5-hydroxymethylcytosine (hmC) (14, 15). This modified nucleobase was initially considered just an intermediate step in the de-methylation pathway (5-Methyl→ hydroxymethyl→ formyl→ carboxyl→ CO_2_) (16, 17), but different studies on the expression of the protein catalyzing mC→hmC conversion (Tet: Ten-Eleven translocation family of dioxygenases)(18–22) and genome wide analysis have shown a crucial role of hmC in the transcription (23–25), especially in processes related to tissue differentiation (26–28). Interestingly, for still unknown reasons, hmC is prevalent in specific brain cell types, as well as in embryonic stem cells (ESCs), with consequential reduction during differentiation (29–33). When present, hmC signals are not randomly distributed, but appear located near transcriptional starting sites and promoters, as well as in certain CpG islands (24, 34–36), suggesting a specific role of this epigenetic modification in gene activity.

The presence of hmC seems to correlate with a reversion of mC effects (37), but the molecular mechanisms connecting the presence of the hmC with gene activity are unclear. Two main possibilities emerge: i) a change in the DNA physical properties reverting to the unmethylated situation that leads to nucleosome repositioning; ii) a direct protein-mediated mechanism. Using a variety of experimental and theoretical methods, we explore here both possibilities. We found that physical properties of hmC are significantly different to those of unmodified cytosine, but not dramatically different to those of mC, suggesting that the reversion of methylation effects in gene expression by TET action cannot be explained simply by change in physical properties of DNA involving nucleosome remodelling. No specific hmC-binding proteins have been described, to our knowledge, but the analysis of known MBDs (16, 38) show a surprisingly wide range of substrate specificity. Our calculations suggest MBD with dual activation/repression activity shows nearly equal affinity for mC and hmC, while those with exclusively inactivating role have a large preference for the methylated form. This can be the ultimate reason of the differential effect of mC and hmC in gene expression.

## MATERIALS AND METHODS

### UV-monitored thermal-denaturation studies

Modified and unmodified 8-nucleotide DNAs (CGAC*GTCG) were synthetized (details in Supporting Methods, Supplementary Table S1) and their absorbance versus temperature curves of each duplex was measured at 5 μM, 20 μM and 66 μM strand concentration in phosphate buffer (25 mM) containing NaCl (100 mM). Experiments were performed in Teflon-stoppered quartz cells of 1 cm and 1mm length path on a JASCO V-650 spectrophotometer equipped with thermoprogrammer. The samples were heated to 90 °C, allowed to cool slowly to 25 °C, and then warmed during the denaturation experiments at a rate of 0.5 °C min^−1^ to 90 °C. The absorbance was monitored at 260 nm. The data were analyzed by the denaturation-curve processing program, MeltWin v.3.0. Melting temperatures (T_m_) were determined by computer fit of the first derivative of absorbance with respect to 1/T. These values were designed to give uniformly separated data points on ln (C_t_/4) scale. ΔH, ΔS, and ΔG values were estimated from dependence of melting temperatures on DNA concentrations (39). The reciprocal values of average melting temperatures were plotted against ln (C_t_/4) and fitted to linear relationships, 1/T_m_ = (R/ΔH) ln(CT/4) + (ΔS/ΔH).

### NMR

Samples of the duplexes with a central CpG step: d(CGTC*GACG)·d(CGTC*GACG) and d(CGAC*GTCG)·d(CGAC*GTCG) with and without epigenetic modifications (synthesis details in Supplementary Methods) were dissolved in either D_2_O or 9:1 H_2_O/D_2_O in 100 mM NaCl and 10 mM sodium phosphate, pH=7 buffer. All NMR spectra were acquired in Bruker Avance spectrometers operating at 600 and 800 MHz, equipped with cryoprobes and processed with the TOPSPIN software. DQF-COSY, TOCSY and NOESY experiments were recorded in D_2_O at 25°C. In the experiments in D_2_O, presaturation was used to suppress the residual H_2_O signal. The NOESY spectra in D_2_O were acquired with a mixing time of 250 ms. TOCSY spectra were recorded with standard MLEV17 spinlock sequence, and a mixing time of 80 ms. NOESY spectra in H_2_O were acquired with 150 ms mixing time at 5°C to reduce exchange with the water. In 2D experiments in H_2_O, water suppression was achieved by including a WATERGATE module prior to acquisition. The spectral analysis program SPARKY (D. K. TD Goddard and D. G. Kneller, SPARKY v3, University of California, San Francisco) was used for semiautomatic assignment of the NOESY cross-peaks and quantitative evaluation of the NOE intensities. Additional details of the NMR analysis are given in Supplementary Material.

### Refinement of the experimental NMR structures

Ensembles of structures using atomistic force-fields based on the distance constraints obtained experimentally were calculated using classical and usual annealing procedure similar to that used to refine most NMR structures (40). Accordingly, ideal fiber B-DNA and A-DNA structures are thermalized (298 K) and equilibrated for 100 ps each (using the same options described previously), applying harmonic restraints of 100 kcal/mol·Å^2^ on the DNA. Then, a 500 ps MD simulation is performed where the global restraints are replaced by the specific NMR distance constraints obtained experimentally. To obtain the final ensemble, 50 structures were chosen and minimized *in vacuo*, keeping the NMR constraints.

### HydroxyMethylCytosine Parameterization

Geometrical parameters for hmC were derived starting from parmbsc1 (41) cytosine and serine from ff14SB for hydroxymethyl group. Final hmC geometry was obtained by energy minimization (42). Charges were derived from RESP fit at HF/6-31G* level starting from an optimized hmC capped at the C1’ using the procedure described in reference (43). Library with parameters for hmC are available upon request.

### Molecular dynamics simulations

We performed molecular dynamics (MD) simulations for 33 12-mer DNA duplexes of sequence CGCGXC*GYCGCG, being C*= C, mC or hmC, with X and Y selected to represent all the 10 possible unique tetranucleotide environments containing a central CG step, were each simulated for 0.5 μs. Additionally the poli(CpG) sequence C*GC*GC*GC*GC*GC*G being C*= C, mC or hmC were simulated. Starting structures were taken from Arnott canonical B-DNA (44), and oligomers were built using the Nucleic Acid Builder (45), epigenetic modification (mC and hmC) were afterwards added to the duplex. The oligomers were simulated using periodic boundary conditions with a truncated octahedral box and an explicit solvent of TIP3P water molecules (46), with a minimum thickness of 10 Å around the solute. The DNA net charge was neutralized with K^+^ cations and K^+^Cl^−^ ion pairs substituted water molecules to reach a concentration of ~0.15 M. Each oligomer was simulated using the AMBER 18 (47) program suite and considering the 3 epigenetic modifications led to a total of 33 simulations and 16.5 μs of unrestrained trajectories. The simulation was performed using our well-established multistep protocol (48–50) which involves energy minimizations of the solvent, slow thermalization and a final re-equilibration for 10 ns, prior to the 0.5 μs production runs. All simulations were carried out in the isothermal-isobaric ensemble (T = 298 K, P = 1 atm) using the parmbsc1 force field (41) and Dang parameters for ions (51, 52). Long-range electrostatic effects were treated using the Particle Mesh Ewald method (53) using a cutoff of 10 Å. The bonds involving hydrogen were restrained using SHAKE (54) and The temperature and the pressure, with a coupling constant of 5 ps were controlled using Berendsen algorithm (55). Trajectories were stored in our BigNAsim database (56), following recent recommendations for FAIR trajectory storage (57).

### Analysis of the trajectories

All the trajectories were processed with the cpptraj module of the AmberTools 18 package (47). The ability of DNA to recognize sodium was analysed using our classical molecular interaction potential (cMIP (58)). The electrostatic interaction term was determined by solving the linear Poisson–Boltzmann equation, while the van der Waals contribution was determined using standard AMBER Lennard–Jones parameters. The ionic strength and the reaction-field dielectric constant were set to 0.15 and 78.4 M, respectively, while the dielectric constant for DNA was set to 8 (59). The molecular plots were generated using either VMD 1.9 (60) or the UCSF Chimera package version 1.8.1 (61).

### Analysis of DNA physical properties and Deformation Energy calculation

DNA helical parameters and backbone torsion angles were measured with the Curves+ and Canal programs (62) along the MD simulations and for X-ray crystal structures. From the ensemble of MD simulations, the covariance matrix describing the deformability of the helical parameters for each dinucleotide step was computed and inverted to generate the 6×6 stiffness matrix, resulting in the force constants for helical deformation at the base pair step level (63, 64). Average and force constants were calculated for the last 200ns of each MD simulation. The position of each cation in curvilinear cylindrical coordinates for each snapshot of the simulations with respect to the instantaneous helical axis was determined using Canion, a utility program that reads the ion counts (65). Deformation energy for the wild type, mCpG and hmCpG DNA sequences was calculated using our mesoscopic energy model (13), with parameters fitted (see above) for each epigenetic modification in each tetramer environment (see above).

### Circularization Assays

#### Phosphorylation and radioactive labeling

Every single stranded oligonucleotide (1 nmol) was 5’-end-phosphorylated with 40U of T4 Polynucleotide kinase (New England Biolabs) by incubation at 37°C in 1x T4 DNA Ligase Reaction Buffer.

#### Annealing and ligation

The complementary strands were denatured at 90°C and subsequently annealed by gradually decreasing the temperature to room conditions. Double stranded oligonucleotides were then multimerized with 400U of T4 DNA ligase (New England Biolabs) by an overnight incubation at room temperature. The reaction was stopped by inactivating the ligase at 65°C for 10 minutes. The DNA was ethanol precipitated and dissolved in 10 μl of DNAse free water.

#### Two-Dimensional Gel Electrophoresis

The ligation/digestion products were loaded on a 5% polyacrylamide native gel (19:1 acrylamide:bisacrylamide) and electrophoresed at room temperature at 4 V/cm in TBE buffer. After separating the different DNA species on the first dimension, the lanes of interest were excised from the gel and loaded on a two dimensional gel (8% polyacrylamide) to separate circular and linear DNA. The separation of the two families of DNA was facilitated by the presence of chloroquine phosphate (250 μg/ml) that distorts the linear molecules of DNA in a different way from the circular ones (3). The gels were stained with *SyBr Safe* (Invitrogen) and visualized on the *ImageQuant* system (GE Healthcare) under UV light (see Results).

### *In vitro* nucleosome reconstitution

Free energies for histone binding in *in vitro* nucleosome reconstitution were measured for the wild type 601 sequence and the epigenetically mutate versions, mCpG601 and hmCpG601 (details in Supplementary Methods). The 3 forms of the high affinity 601.2 DNA sequence (sequences in Supplementary Information), with normal C-, mC-, hmC-DNA were chemically synthetized starting from the oligos 65-, 66-, 80- and 81-nucleotide long DNAs on the 0.2 μmol scale on an Applied Biosystems 394 synthesizer (see Supplementary Methods for details, Supplementary Table S1). We introduced 2 CpG base pair steps 5 or 11 base pairs apart: in positions that face the protein core through the minor and major groove or, alternatively, only the major grooves. Such positions explore sites in the nucleosomal DNA of negative and positive opening of the base pairs along their long axis (13). Bands intensities corresponding to nucleosome-bound or free DNA were measured by densitometry using the *PhosphorImager* system (GE Healthcare) (Supplementary Figure S1). Bands intensities quantitation allowed to calculate the affinity for each sequence to form nucleosome as the ratio of counts in nucleosome bound DNA to counts in free DNA. Experimental normalised affinity was then computed dividing the relative affinity for methylated or hydroxymethylated sequences by the control (unmodified cytosine). Each experiment, for each sequence, was repeated four times.

### Thermodynamic Integration

We used a thermodynamic cycle to compute the reversible work associated to the alchemical transformation between two DNA sequences (between mC and hmC), both in the bound and in the unbound state for a list of all MDB protein-DNA complexes (Supplementary Methods). The details on how these calculations were carried out are explained in a previous work in which we used the same method to calculate the variation in free energy adding methylation in a nucleosome complex (13). In this case, to achieve the alchemical transformation between mC and hmC, our molecular dynamics simulations convert the hydroxyl group in position 5 of the cytosine ring into a hydrogen atom. In each complex for each mutation, the variation of λ is discretized in 21 windows, i.e. dλ=0.05, and the final ΔG is computed via numerical integration. We used soft-core potentials implemented in GROMACS to avoid singularities in the Lennard-Jones and Coulomb potentials, with α=0.3, σ=0.25 and a soft-core power of 1. The initial structures for each window were obtained from a 10 ns simulation in which λ was continuously varied from 0 to 1. From this simulation we extracted frames corresponding to a given λ value, and we relaxed the structures by minimizing the energy of the system in those configurations. We simulated each window at fixed λ value for 1 ns, and we discarded the first 100 ps of simulation. For each window we collected 9 estimates for 〈(δH)/δλ〉_λ by using 9 blocks of 100 ps, which were then integrated through the entire mutation pathway to obtain mutation free energies (with associated statistical errors).

## RESULTS AND DISCUSSION

### The effect of 5-hydroxymethylcitosine on the stability of DNA

We first analysed the changes in stability due to methylation and hydroxymethylation of the model sequence d(CGAC*GTCG)·d(CGAC*GTCG). UV-Melting experiments demonstrated (see Methods and Supplementary Table S2) a slight increase in the melting temperature of hmC with respect to unmodified DNA (2-3 degrees), and a slight decrease (around 1 degree) from mC-DNA. Processing of melting profiles confirms the small impact of C→mC→hmC substitutions in duplex stability, and the predictable enthalpy/entropy compensation, with the hydrophobicity term favouring (as expected, (66)) the stability of mC (mC >hmC>C, see Supplementary Table S3).

### The effect of 5-hydroxymethylcitosine on dsDNA structure. NMR studies

We studied the impact of hmC at d(CpG) steps on the conformation of DNA by NMR experiments using two model sequences with a central CpG step: d(CGTC*GACG)·d(CGTC*GACG) (central base pair very flexible, (67)) and d(CGCGAC*GTCGCG)·d(CGCGAC*GTCGCG) (central base pair very stiff, (67)). NMR data was collected for C*=C, hmC and C*=C, mC and hmC, respectively; allowing us to obtain, for the first time direct experimental information on the structure of DNA related to epigenetic changes C←→mC←→hmC placed on both strands of the duplex (i.e., d(C*pG)·d(C*pG)) (see Methods and Supplementary information). Sequential assignments of exchangeable and non-exchangeable proton found small, but significant changes in chemical shifts of the groups involved in H-bonding at the substitution site (chemical shifts at C* amino move from 8.21 ppm (C*=C) to 8.41 ppm (C*=mC) and 8.54 ppm (C*=hmC)) (data in Supplementary Tables S4-S9 and Figures S2-S5). The NMR signal of −OH in the hmC duplex could not be detected, most probably due to rapid exchange with the solvent, suggesting that OH-N6 hydrogen bond is not highly populated. Angular and distance constraints derived from J-couplings and NOE cross-peaks, respectively, allowed us to determine the solution structures (see Methods and Supplementary information) for the mC and hmC containing duplexes in the two distinct tetrameric environments. In all cases structures agree very well with a general B-like duplex (Figure 1). The changes induced by the presence of mC or hmC are small, in general less important than those induced by tetramer-dependency (TC*GA vs AC*GT) and a detailed analysis at the central base pair step (d(CpG); see Table 1) demonstrates that at the very local level the structural impact of epigenetic changes in DNA are modest.

**Figure 1.**
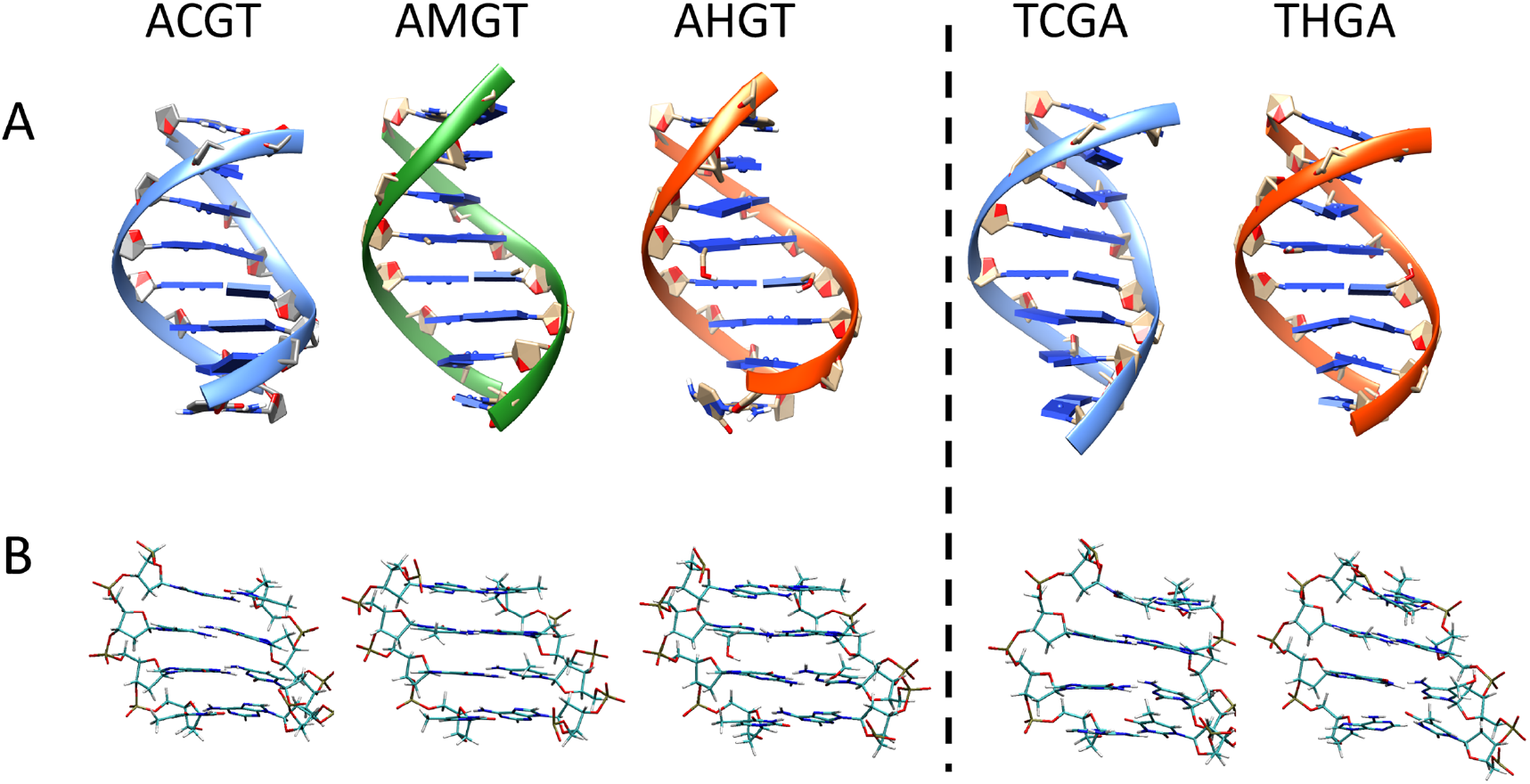
Average NMR structure of the sequences d(CGTC*GACG)·d(CGTC*GACG) and d(CGCGAC*GTCGCG)·d(CGCGAC*GTCGCG) (C* being C, mC or hmC respectively) for the central octamer (A) and the central tetramer (B).

**Table 1.**
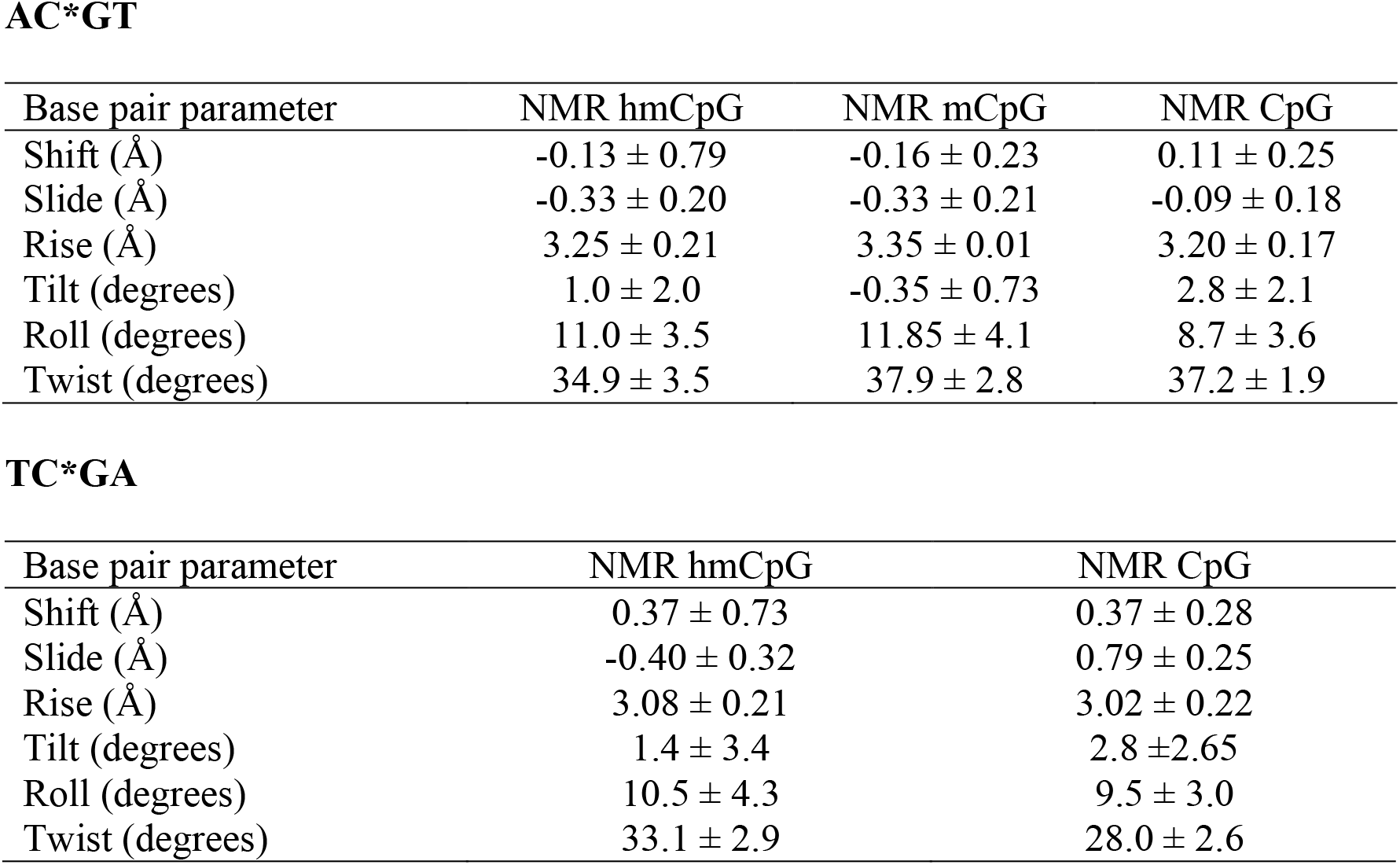
Average parameters (in Å and Degrees) with standard deviation for the central step in the NMR structure of the sequences d(CGTC*GACG)·d(CGTC*GACG) and d(CGCGAC*GTCGCG)·d(CGCGAC*GTCGCG) (C* being C, mC or hmC).

AC\*GT
| Base pair parameter | NMR hmCpG | NMR mCpG | NMR CpG |
| --- | --- | --- | --- |
| Shift (Å) | $-0.13 \pm 0.79$ | $-0.16 \pm 0.23$ | $0.11 \pm 0.25$ |
| Slide (Å) | $-0.33 \pm 0.20$ | $-0.33 \pm 0.21$ | $-0.09 \pm 0.18$ |
| Rise (Å) | $3.25 \pm 0.21$ | $3.35 \pm 0.01$ | $3.20 \pm 0.17$ |
| Tilt (degrees) | $1.0 \pm 2.0$ | $-0.35 \pm 0.73$ | $2.8 \pm 2.1$ |
| Roll (degrees) | $11.0 \pm 3.5$ | $11.85 \pm 4.1$ | $8.7 \pm 3.6$ |
| Twist (degrees) | $34.9 \pm 3.5$ | $37.9 \pm 2.8$ | $37.2 \pm 1.9$ |

| Base pair parameter | NMR hmCpG | NMR CpG |
| --- | --- | --- |
| Shift (Å) | $0.37 \pm 0.73$ | $0.37 \pm 0.28$ |
| Slide (Å) | $-0.40 \pm 0.32$ | $0.79 \pm 0.25$ |
| Rise (Å) | $3.08 \pm 0.21$ | $3.02 \pm 0.22$ |
| Tilt (degrees) | $1.4 \pm 3.4$ | $2.8 \pm 2.65$ |
| Roll (degrees) | $10.5 \pm 4.3$ | $9.5 \pm 3.0$ |
| Twist (degrees) | $33.1 \pm 2.9$ | $28.0 \pm 2.6$ |

### The effect of 5-hydroxymethylcytosine on the physical properties of DNA

Unbiased MD ensembles of d(CGTC*GACG)·d(CGTC*GACG) and d(CGAC*GTCG)·d(CGAC*GTCG) agrees very well with the MD-NMR-biased ones (RMSd deviation around 0.7-1.1 Å from the average NMR structure for the central tetramers; see detailed analysis at the C*pG step in Supplementary Figure S6), supporting the quality of parmbsc1 simulations (41). This allows us to analyse the impact of C→mC→hmC substitutions in a much wider range of sequence environments and in the absence of the artificial restraints used in NMR refinements. Such restrains which are nearly-harmonic bias the intrinsic deformability of the DNA hindering the possibility to sample for example, the bimodality at the central CpG step, a well-known polymorphism found both in MD simulation and in high-resolution experimental structures (67–70). With this aim we collected 0.5 μs trajectories (see Methods) of 10 dodecamers containing all the unique tetramer environments of the d(C*pG) step (for C*=C, mC or hmC). Additional trajectories were collected for d(C*pG)_6_·d(C*pG)_6_, a poly(CpG) dodecamer mimicking CpG islands, repetitive polymers with unique biological importance. In summary, we collected 16.5 μs of unbiased trajectories of dodecamers containing C*pG tetramers (0.5 μs x 33 structures), which confirm that the introduction of epigenetic variants of cytosine leads to only mild structural alterations in the duplex, and the differences between mC- and hmC-DNAs being very small (Supplementary Table S10). Modifications induced by epigenetic changes are typically smaller than tetramer-induced changes, but there is a clear general tendency for a reduction in twist and an increase in roll when C is substituted by mC or hmC (Figure 2).

**Figure 2.**
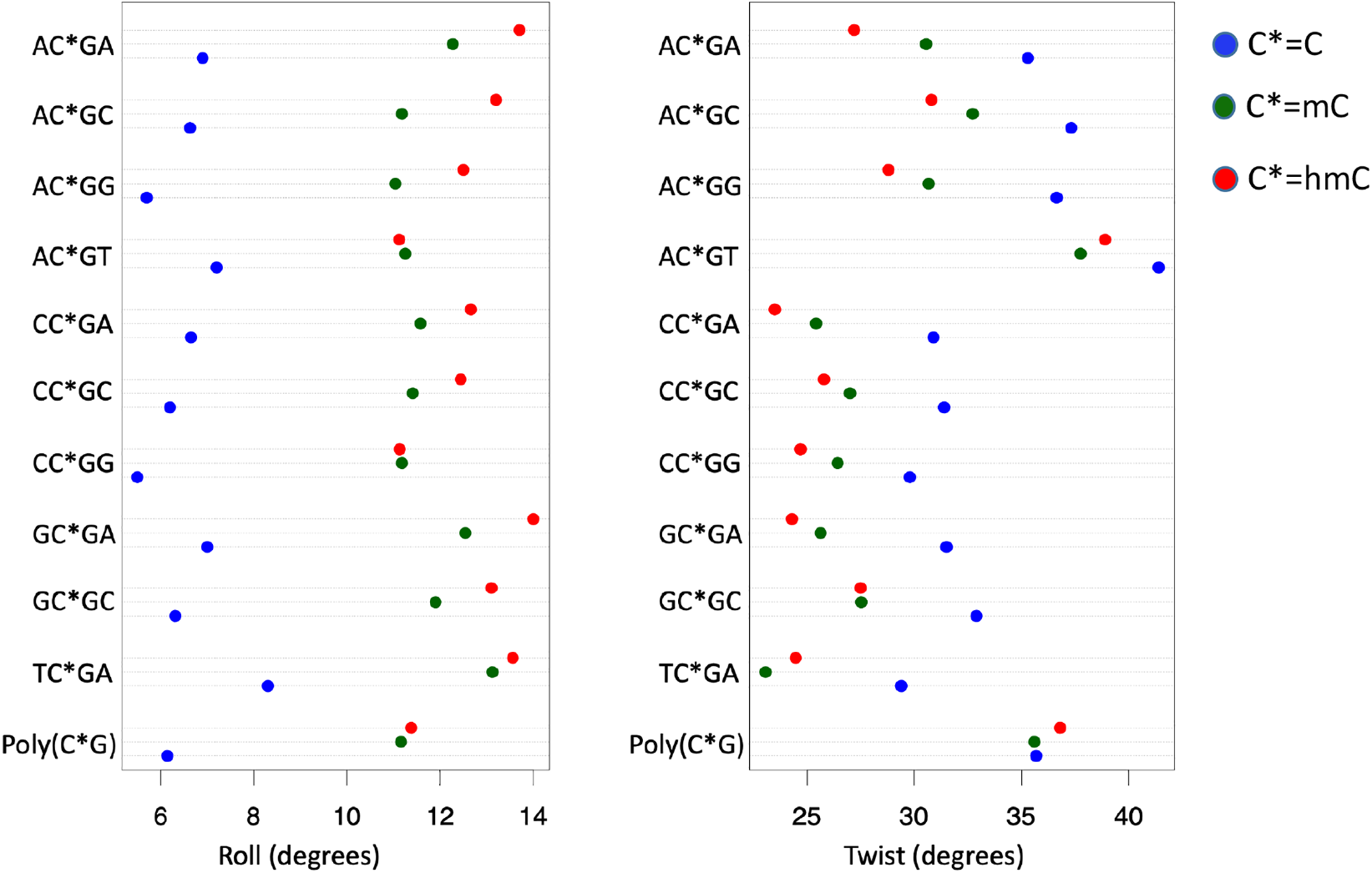
Average value for the base pair parameters roll (left panel) and twist (right) for C*G step in the different tetramer environment. Being C* normal cytosine (blue dots), methylated cytosine (green dots) and hydroxymethylated cytosine (red dots).

In the case of twist, the epigenetic changes induce (in general) a displacement of the bimodal CpG distribution towards low twist values (Figure 3), while in the case of roll, it results in the movement of the normal distribution towards higher values, probably related to the steric hindrance of the cytosine 5-substituent (Figure 3). Note that it is difficult, due to the refinement procedures, to detect changes in twist bimodality in NMR structures, but the high twist to low twist transition detected by MD in flexible sequences has been already detected in a high-resolution crystal structure (PDB ID 4GLH, 4HLI and 4GLC) (see Supplementary Figure S7), supporting the validity of our theoretical findings. Finally, and very interestingly, we found that some tetramers show quite unique distributions (for example the high twist of AC*GT) and that polymer effects are not negligible as shown in the d(CGCG)·d(CGCG) tetramer behaves differently when embedded in the reference dodecamer or in a poly(CpG).

**Figure 3.**
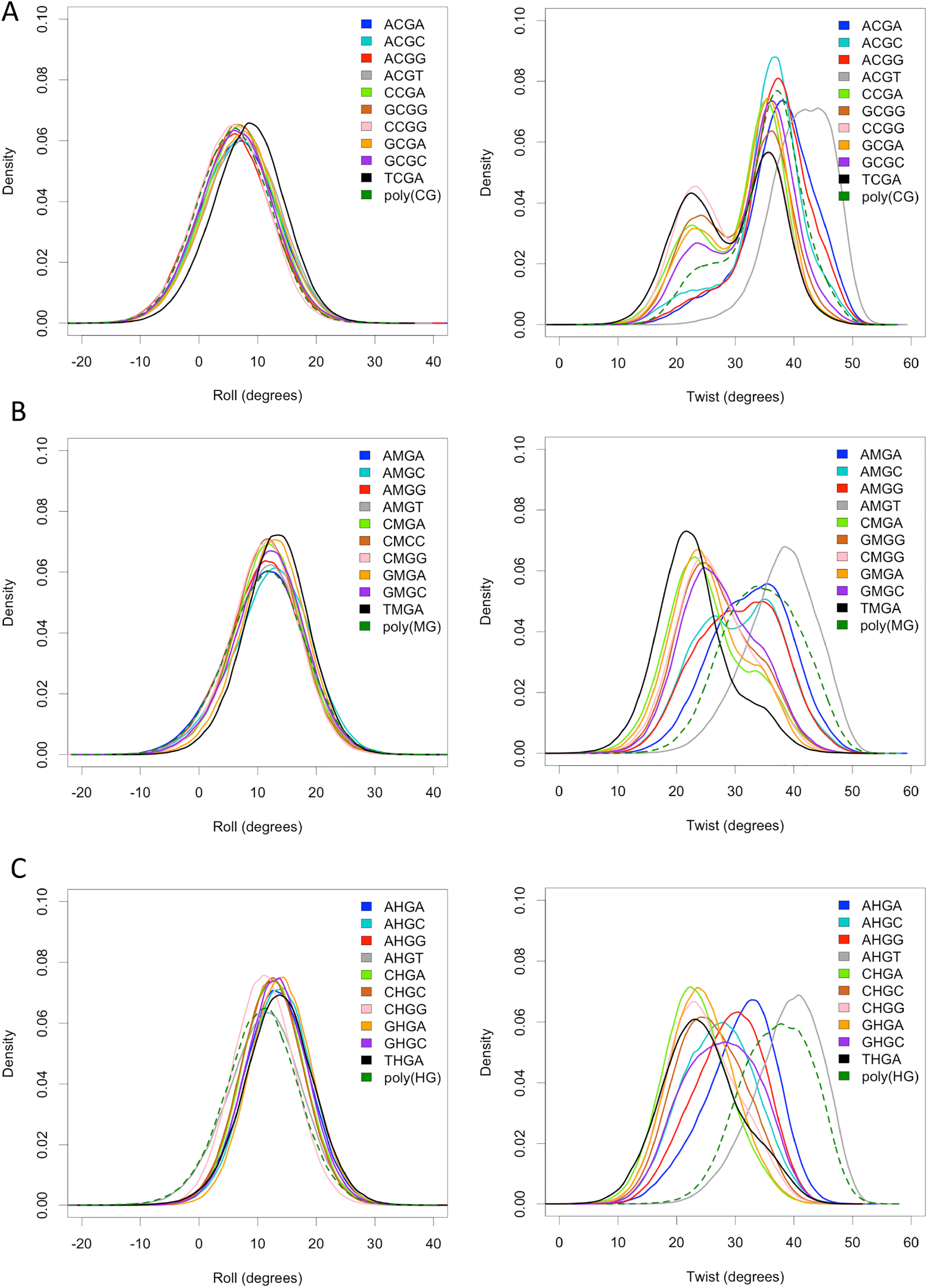
Roll (left panel) and twist (right panel) distributions for each C*pG step, in a different tetramer environment, being C* normal (A), methylated (B) and hydroxymethylated (C). Dashed line for C*pG step in a poly(CpG) sequence.

Stiffness analysis (see Methods) (63, 64) illustrates the impact of epigenetic changes on the elastic deformability of DNA. In general, cytosine methylation increases the stiffness of CpG steps, mainly due to the restriction of roll flexibility. Such a stiffness increase is even more evident for hmC (see Table 2 and Supplementary Table S11), due the narrowing of the roll distribution (see Figure 3), probably related to the formation of transient hydrogen bond interactions involving the hydroxyl group of hmC (see Supplementary Figure S8 as examples). It is worth noting, again, that these general trends are not universal and significant deviations are evident for certain tetramers (for example ACGT and TCGA). Again poly(CpG) has a rather differential behaviour, as for this repetitive sequence methylation increases rather than decrease flexibility, which combined with results above suggest that epigenetic changes can affect physical properties of CpG islands in a quite different way than in the rest of the DNA. This unique behaviour may explain claims by other authors of a general increase in DNA flexibility induced by cytosine methylation (71).

**Table 2.**
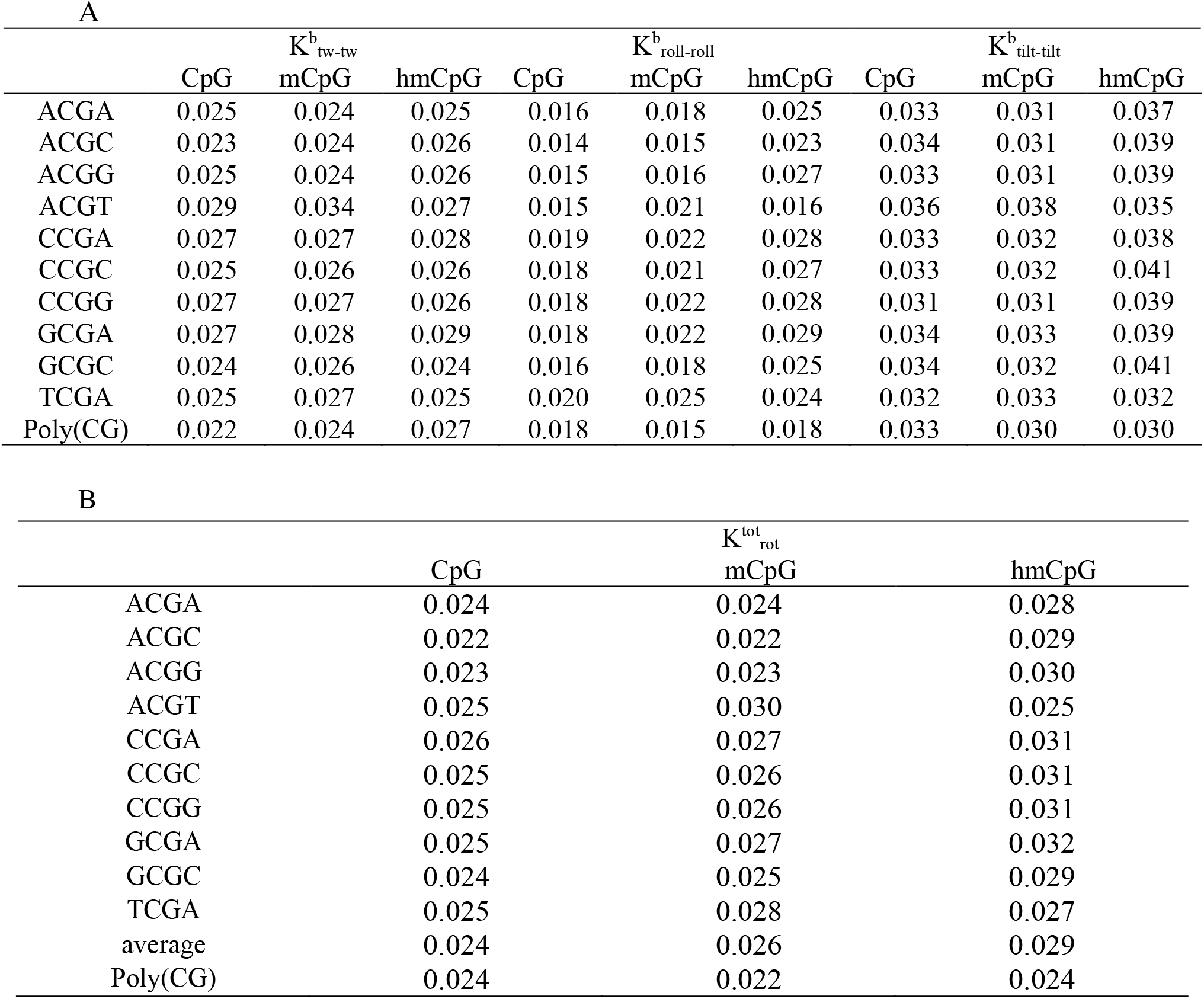
A) Diagonal stiffness constants for rotational movements in kcal/mol deg^2^ for the central C*pG step (C*=C, mC and hmC) in the different tetrameric environments. B) Cubed root of the product of the diagonal stiffness constants for rotational movements in kcal/mol deg^2^.

To validate our claim that epigenetic variants tend to increase the DNA stiffness (in the majority of the sequence contexts), we carried out DNA circularization experiments using the double stranded 21mers (d(GAAAAAACGGG**C*G**AAAAACGG)· d(TCCCGTTTTT**C*G**CCCGTTTTT)) with a central C*pG dinucleotide (C*=C, mC, hmC, respectively) as described in Perez *et al* (12). Under favorable ligation conditions (see Methods), the multimers form circles that are as short as allowed by the flexibility of the DNA. Results shown in Figure 4 clearly demonstrate that, as suggested by our MD simulations (Table 2), epigenetic variants increase (in most sequence context) the stiffness of DNA, following the order: hmC>mC>C. In summary, the reversion of methylation signature by hydroxymethylation cannot be explained as a recovery of the properties of the unmethylated DNA.

**Figure 4.**
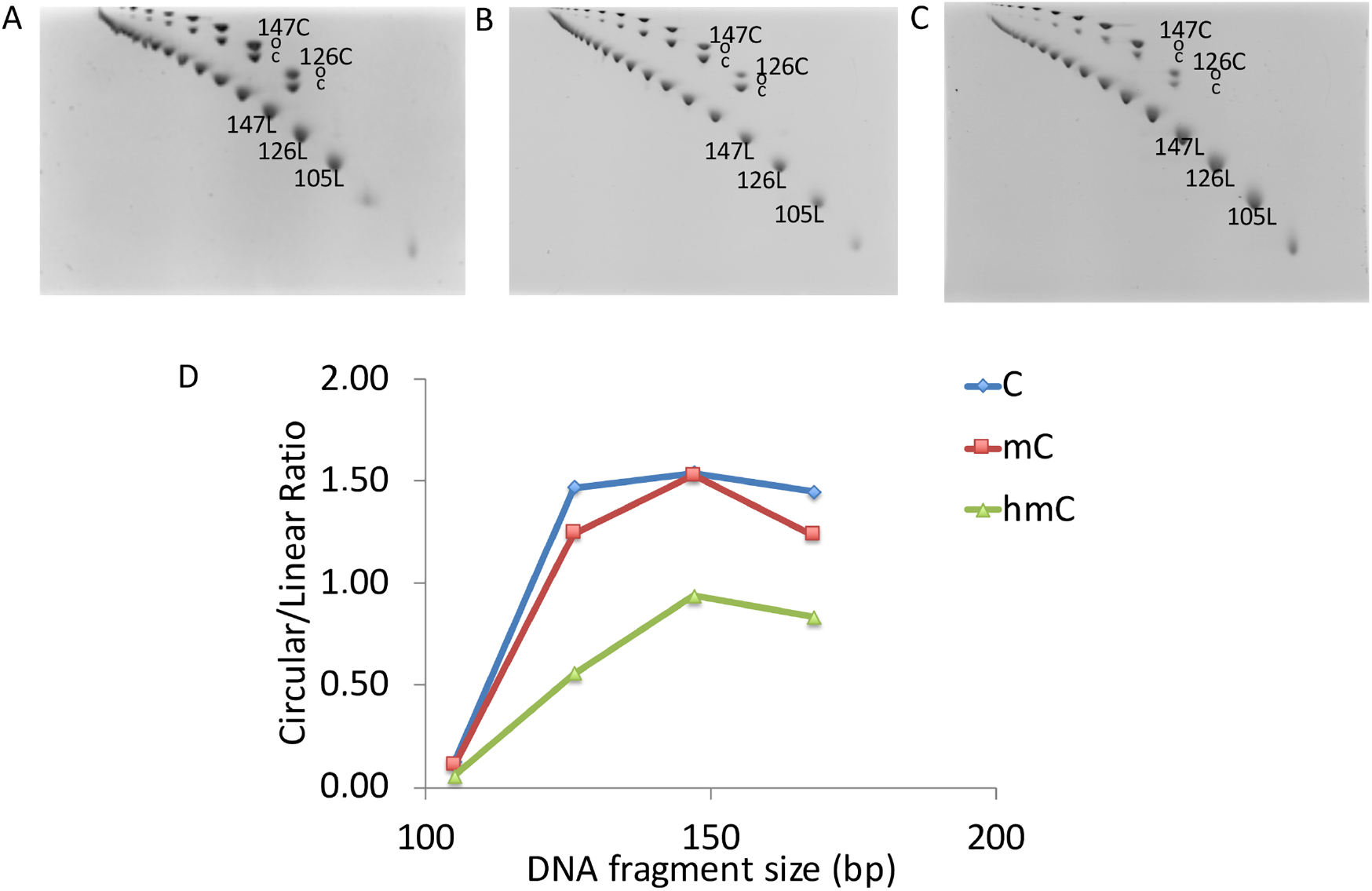
(A-C) 2D polyacrylamide native gels showing different migrations of linear and circular DNA species oligomers of 21 bp, respectively for (A) Cytosine (C), (B) Methylcytosine (mC) and (C) Hydroxymethylcytosine (hmC) containing fragments. Linear DNA molecules are positioned on the lower diagonal, and circular DNA molecules are positioned on the upper diagonal. (D) The circularization efficiency expressed as the ratio between Circular and Linear molecules of the same size (base pairs, bp).

The reduction of flexibility introduced by epigenetic variants of cytosine is expected to increase the energetic cost of wrapping the DNA around the nucleosome core (13), leading to a decrease in nucleosome affinity. To confirm this hypothesis, we compute deformation energies (see Methods) associated to nucleosome wrapping for DNA containing unmodified CpG, mCpG and hmCpG steps. To include phasing issues into consideration we explored two scenarios: i) 4 epigenetic changes (2 d(CpG) steps modified) “in-phase”, i.e., each modification separated by 11 bases (the periodicity of DNA in a nucleosome (72)) and ii) 4 epigenetic changes in “anti-phase”, i.e. each modification separated by 5 bps. Results in Figure 5A demonstrate that epigenetic variations increase the energy cost required for nucleosome wrapping, both when introduced in “in-phase” and “anti-phase” (Supplementary Figure S9), suggesting that change in flexibility rather than in the intrinsic curvature (roll and tilt) are the main responsible of the epigenetic-induced difference in nucleosome affinity. To validate these theoretical findings, we performed nucleosome reconstitution experiments (see Methods, Supplementary Methods) (73) using both “in-phase” and “anti-phase” arrangements (the same sequences as before). Results (Figure 5B) confirm that the C→mC→hmC substitution leads to a reduction in the efficiency of nucleosome reconstitution, the difference between methylated and hydroxymethylated DNAs being small. Interestingly, experimental changes in nucleosome reconstitution induced by epigenetic alterations “in-phase” are not dramatically different than those found in “anti-phase”, where the effect of changes in intrinsic curvature should cancel, supporting the claims that changes in flexibility are the main responsible for epigenetic-related changes in nucleosome affinity.

**Figure 5.**
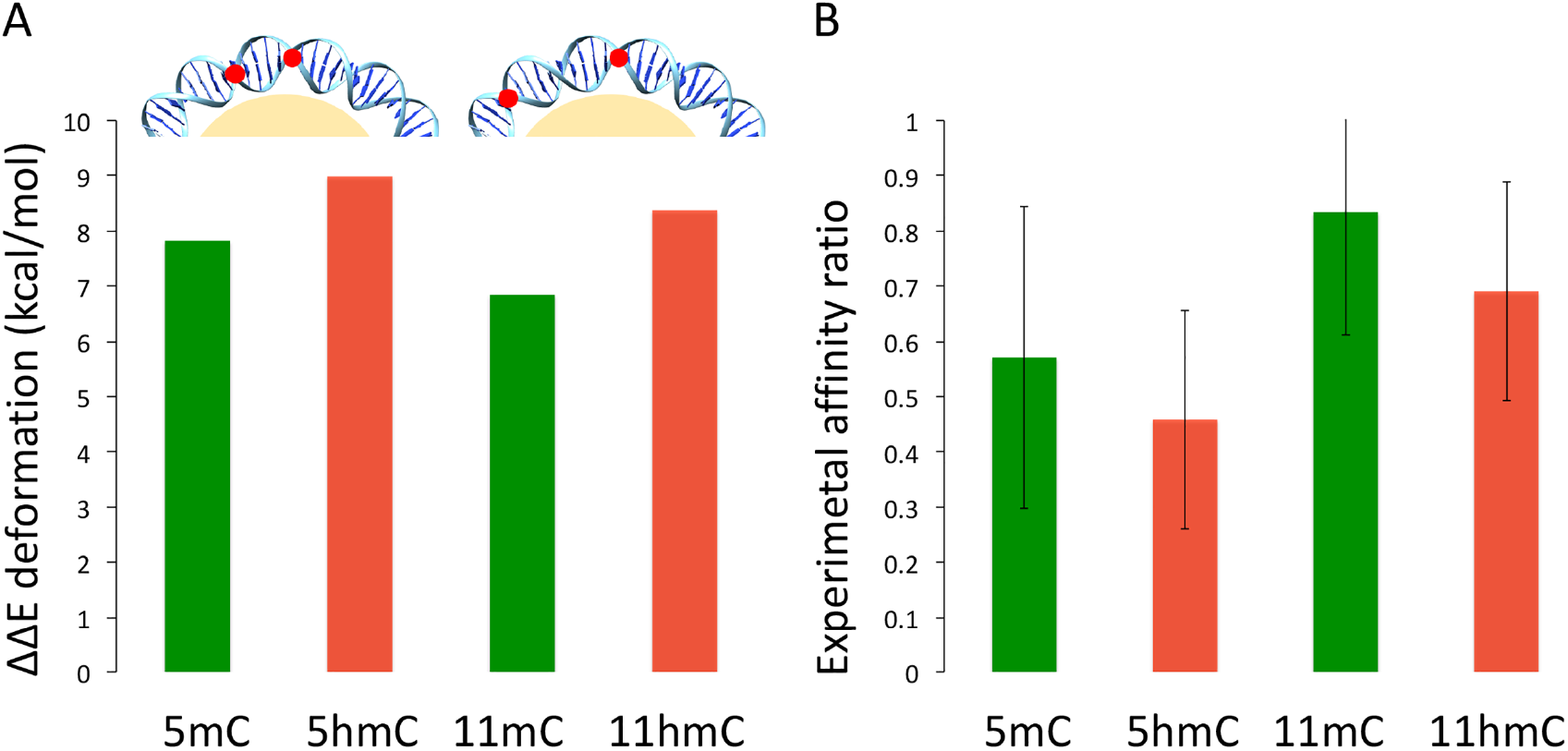
A) Deformation energy variation cost required for nucleosome wrapping, difference between the energy cost of the methylated (mC) and hydroxymethylated (hmC) DNA respect to the control (unmodified cytosine). At the top a schematic view of the positioning of the modifications (red dots), respectively 5 and 11 base pairs apart on the DNA wrapped around the histones (in yellow). B) Experimental affinity ratio, with standard deviation, between the methylated (mC) and hydroxymethylated (hmC) DNA respect to the control (unmodified cytosine) calculated using nucleosome reconstitution

Altogether, theoretical and experimental results strongly suggest that C→mC→hmC substitution at d(CpG) steps leads to small changes in structure and to a small, and environment specific increase in the stiffness (hmC>mC>C), which is reflected in a reduction of nucleosome affinity. Very interestingly, the behavior of DNA containing hmC is much closer to that of met-DNA than that of unmodified DNA, which means that the “reversion” to the unmodified DNA behavior found when methylated DNA is processed by TET-proteins to yield hmC-DNA (22) cannot be explained by changes in the intrinsic physical properties of DNA and the associated recovering of the nucleosome distribution.

### The impact of epigenetic modifications on the recognition properties of DNA

The preferred recognition pattern of DNA changes with epigenetic modifications and this can modulate DNA-protein contacts. Thus, DNA containing CpG steps exhibits a preferred region of interaction with cations along the minor groove (Figure 6) (see Methods) (65). Cytosine methylation widens slightly the minor groove, displacing the preferred interaction region to the backbone and hydroxymethylation makes the major groove the preferred interaction region (see examples in Supplementary Figure S10), probably as a consequence of the presence of a polar group. The polarity of the CH_2_OH group is also visible in the water distributions (see examples in Supplementary Figure S11), where the integration of the hydroxyl into highly ordered strings of waters leads in general to a very dense solvation atmosphere along the major groove.

**Figure 6.**
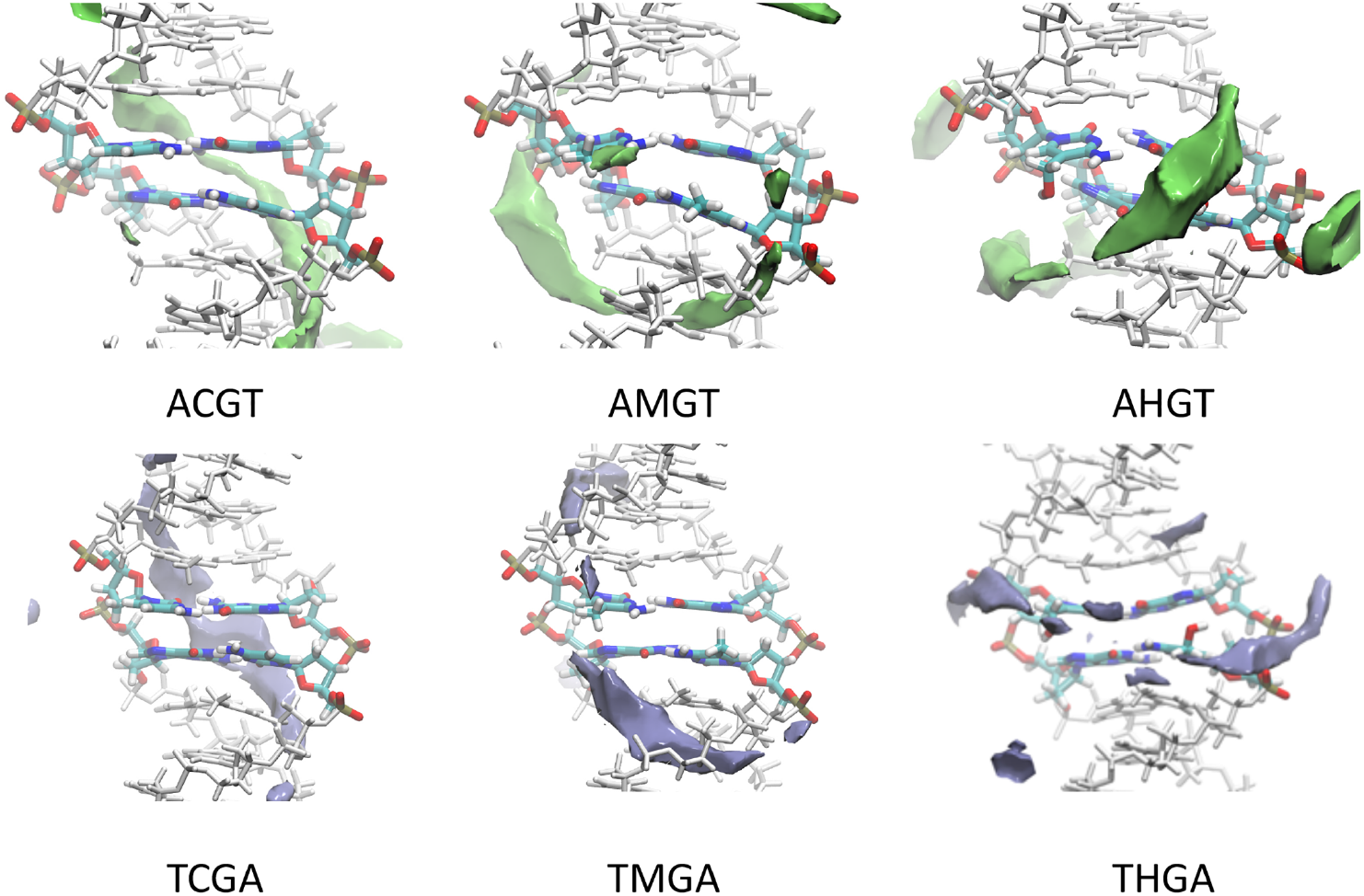
Examples of classical molecular interaction potentials using sodium as a probe (58). For the sake of comparison, all the averaged structure with the 3 epigenetic modifications of the cytosine were aligned and the same isosurface −3.5 kcal/mol was computed.

Results above suggest epigenetic modifications alter DNA interaction pattern, which might impact the binding of methylated DNA binding domains (MBDs). We focus here on the four MBD proteins related to the regulation of gene expression: i) MeCP2, a protein with direct (via repression transcription domain; TRD) and indirect (via recruitment of histone deacetylases) silencing activity in mature nerve cell (3, 38, 74), ii) MBD1 which represses directly (via TRD) or indirectly (via recruitment of histone methylases) gene expression (38), iii) MBD2 which is also coupled to repression (directly through TRD, or indirectly through histone methylation and deacetylation), and which is known to induce NuRD-dependent nucleosome remodeling (38, 75, 76)and finally iv) MBD3 which lacks TRD and which has been related to a myriad of secondary interactions with other partners such as NuRD, DNMT and TET leading to either gene expression or activation.(30, 38, 76).

We built models of the four MBD-DNA complexes from the available experimental data (details in Methods and Supplementary Methods). The complexes were solvated and used as starting conformations for MD simulations, coupled to thermodynamic integration (MD/TI), which combined with standard thermodynamic cycles (Figure 7) allowed us to determine the impact in binding of methyl-cytosine to hydroxymethyl-cytosine change in protein binding (see Methods). Quite interestingly, despite the very high sequence similarity the four studied MBD have a different discriminant hmC vs mC power. Based on our simulations, MBD1 has the highest specificity for mC compared to hmC, MBD2 and MeCP2 show a small preference for the methylated DNA, MBD3 seems unable to distinguish between mC and hmC.

**Figure 7.**
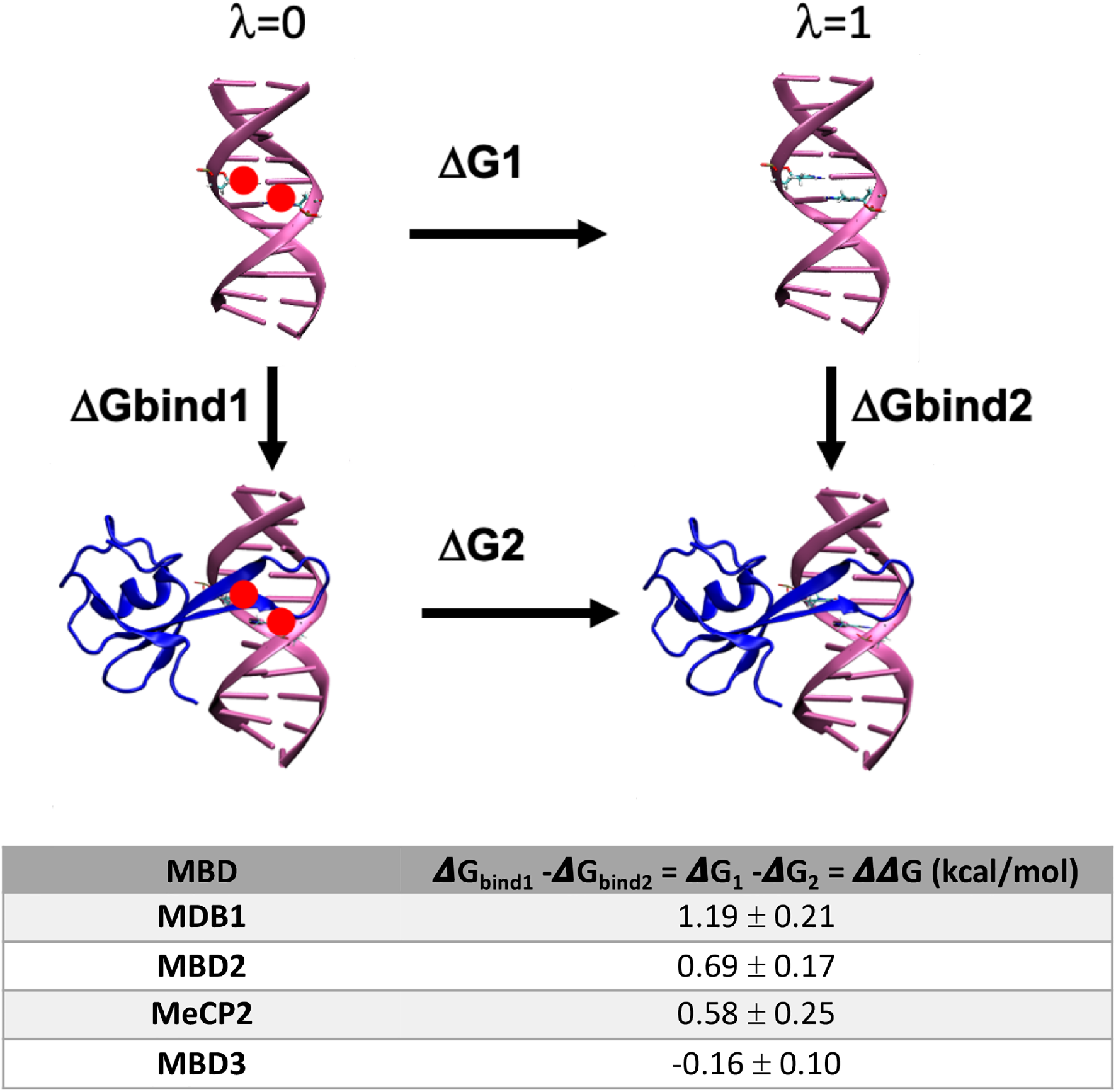
Diagram of the thermodynamic cycle used to extract the free energy variation (DDG (kcal/mol)) in nucleosome-DNA stability due to methylation of CpG steps. The calculations were performed for each different MBD protein-DNA complex, interchanging hmC and mC and computing the associated reversible work by MD/TI. Positive ΔΔG values mean that the complex binding to met-DNA is more favorable than to the hydroxymethylated one.

Analysis of the binding sites (Figure 8) allowed us to determine the origin of the differential specificity. The four proteins have a large sequence similarity and identical fold. They recognize d(CpG) steps (i.e. d(CpG)·d(CpG) by means of interactions along the major groove (N7 and O6) of the two guanines. Such interactions are made by two Arg in three of the proteins (MeCP2, MBD1 and MBD2), while for MBD3 recognition is made by one Arg and one His, suggesting a lower specificity (Figure 8A). The electrostatic environment of the protein pocket where the hydroxyl group is placed is also different, in the MBD1, the protein with highest affinity for mC, the methyl group is placed in a pocket defined by residues such as VAL and PHE defining a quite hydrophobic cavity. When the methyl is hydroxylated, the OH-group is embedded in a unfavourable apolar environment (see white surface in Figure 8B left panel). On the contrary, in MDB3 the hydroxyl group are placed in a hydrophilic/polar cavities, interacting with ASP and a SER (see red and green surface in Figure 8B). These theoretical results are supported by a myriad of experimental evidences, for example, MBD1 is known to be the protein with the largest specificity for methylated DNA, while MBD3 is known to be the most promiscuous of all MBD (38). There is also good agreement with in vitro binding estimates by Hashimoto et al. (77), which suggests the same range of mC/hmC specificity than that found here.

**Figure 8.**
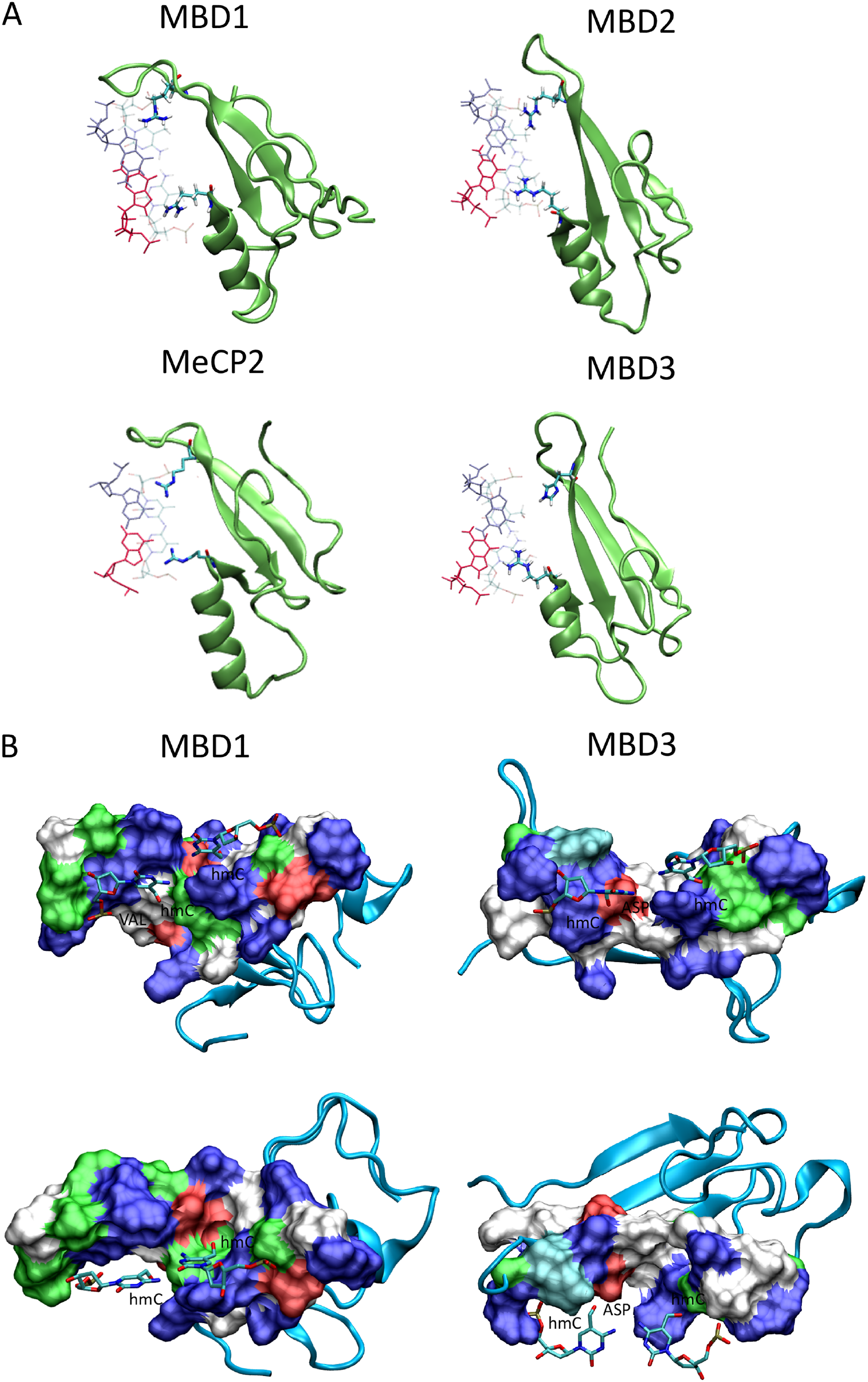
A) Detail of the Arg-recognition model for each MBD-DNA complex (see Methods for detail), arginines interacting with the guanines paired to the epigenetically modified cytosines. B) Protein surface closest to the modified cytosine (hmC), showing the interaction cavity hydroxyl group for the two extreme examples MDB1 (left panel) and MDB3 (right panel).

In summary, MD/TI simulations strongly suggest that mC→hmC introduces a general decrease in binding affinity in MBD proteins with repressive activities, reversing then the inactivating signals associated with cytosine methylation. On the contrary, MBD3, which shows that largest sequence variability at the recognition site has no preference between the two epigenetic variants. This suggests that MBD3-associated signaling is not affected, which might yield to specific activation or deactivation of genes. The final consequences of the presence of d(hmCpG) steps should be then not only a general activation of gene activity (reversing methylation signal), but a qualitative gain in the complex and promiscuous MBD3 associated signaling.

## CONCLUSIONS

The presence of hmC in both strands of a d(CpG) step leads to minor structural changes in the DNA duplex, but to non-negligible changes in physical properties, which are reflected in a general increase in the stiffness of DNA that becomes evident in MD simulations and is validated by circularization experiments. This increase in stiffness implies a reduced ability to wrap around nucleosomes, as suggested by MD simulations and demonstrated by gel reconstitution experiments. Very interestingly, both simulations and experiments suggest that hydroxymethylated DNA is even stiffer than methylated DNA, precluding that physical properties could justify the known reversion to unmethylated behavior which is coupled to the d(mCpG)→d(hmCpG) transformation catalyzed by TET. In fact, simulation and experiments suggest a small impact of d(mCpG)→d(hmCpG) transformation on chromatin conformation.

The presence of the hydroxymethyl in the major groove of changes the recognition properties of DNA and this is expected to modify binding of many effector properties, modulating then in a complex way DNA activation and repression pathways. In this paper we have centered our attention in MBD, the major elements for recognition of methylation signals. Accurate MD/TI calculations strongly suggest that d(mCpG)→d(hmCpG) transformation leads to reduction of the affinity of binding for all MBD coupled to repressive action, while binding to MBD3, the only methyl-binding domain with dual (activation/repression) activities is not affected. Simulations suggest then that a significant part of the biological effect of d(mCpG)→d(hmCpG) transformation can be related to an unbalance in the recognition of DNA by MBD proteins.

## ACKNOWLEDGEMENTS

We thank Irene Gomez-Pinto of the Instituto Química Física Rocasolano. Consejo Superior de Investigaciones Científicas (CSIC) in Madrid for her support in the NMR experiments. NMR experiments were performed in the ‘‘Manuel Rico’’ NMR laboratory (LMR), a node of the Spanish Large-Scale National Facility (ICTS R-LRB).

