## Supplementary Information for "The Impact of The Hydroxymethyl-Cytosine Epigenetic Signature on DNA Structure And Function"

[1]Institute for Research in Biomedicine (IRB Barcelona). The Barcelona Institute of Science and Technology. Baldiri Reixac 10–12, 08028 Barcelona, Spain.
[2]Department of Biological Sciences, CENUR North Riverside, University of the Republic (UdelaR), Gral. Rivera 1350, 50000 Salto, Uruguay.
[3]Functional Genetics Lab., Institute Pasteur of Montevideo, Mataojo 2020, 11400 Montevideo, Uruguay.
[4]Chemistry Department, University of Cambridge, Lensfield Road, Cambridge, CB2 1EW, UK
[5]Instituto Química Física Rocasolano. Consejo Superior de Investigaciones Científicas (CSIC), Serrano 119, 28006 Madrid, Spain.
[6]Department of Biochemistry and Molecular Biology. University of Barcelona, 08028 Barcelona, Spain.

### Supplementary Methods

#### Synthesis of oligonucleotides for NMR studies

mdC-Modified 8mer and unmodified 8mer and 12mer DNAs were purchased from Sigma-Aldrich. hmdC-Modified 8mer and 12mer DNAs were synthesized on the 1 μmol scale using standard phosphoramidite methods (1) (8mer: DMT-ON mode; 12-mer: DMT-OFF mode). Commercially available 5'-*O*-DMT-dG$^{iBu}$-3'-succinyl-LCAA-CPG (Link Technologies) was used as the solid support. Phosphoramidite monomers of dA$^{Bz}$, dC$^{Ac}$, dG$^{iBu}$ and T, and deblocking solution (3% TCA in $CH_2Cl_2$), activator solution (0.3 M 5-benzylthio-1-H-tetrazole in $CH_3CN$), CAP A solution (acetic anhydride/pyridine/THF), CAP B solution (THF/*N*-methylimidazole 84/16) and oxidizing solution (0.02 M iodine in THF/pyridine/water (7:2:1)) were obtained from Link Technologies. 5-Hydroxymethyl-dC$^{Bz}$ CE phosphoramidite was obtained from Glen Research. Except for hmdC phosphoramidite, the standard coupling conditions were used. Coupling time for hmdC was 15 minutes. After solid-phase synthesis, the solid supports were transferred to screw-cap vials and incubated at 75 ºC for 19 h with 1 mL of $NH_3$ solution (33%). After cleavage from the solid support and deprotection, the supernatants were transferred into 2 mL Eppendorf tubes and the supports were rinsed with water (2 x 0.25 mL). The combined solutions were evaporated to dryness using an evaporating centrifuge.

The hmdC-modified 12mer DNA was purified by 20% denaturing polyacrylamide gel; the oligonucleotide was isolated by the crush and soak method and quantified by absorption at 260 nm.

The hmdC-modified 8mer DNA was purified by HPLC (DMT-ON). Column: Nucleosil 120-10 C18 (250 x 4 mm); 20 min linear gradient from 15% to 80% B and 5 min 80% B, flow rate 3 mL/min; solutionA was 5% ACN in 0.1M aqueous triethylammonium acetate (TEAA) and B 70% ACN in 0.1M aqueous TEAA. The pure fractions were combined and evaporated to dryness. The residue that was obtained was treated with 1 mL of 80% AcOH solution and incubated at room temperature for 30 min. The deprotected oligonucleotide was desalted on a NAP-10 column, using water as the eluent, and quantified by absorption at 260 nm.

**Synthesis and preparation of 601, mCpG-601 and hmCpG-601 DNA sequences**

To assess the effect of DNA methylation and its oxidized forms on nucleosome assembly, we selected a nucleosome positioning sequence (DNA construct 601.2 in Anderson and Widom (2)) and synthesized several constructs containing 2 modifications per strand each, separated by 5 or 11 nucleotides (see sequences below - The modified cytosines are shown in bold red).

Position 5: 2 modifications antiphase 5

5'CTGCAGAAGCTTGGTCCCGGGGCCGCTCAATTGGTCGTAGCAAGCTCTGGATC
CGCTTGAT**C**GAA**C**GTACGCGCTGTCCCCCGCGTTTTAACCGCCAAGGGGATTACT
CCCTAGTCTCCAGGCACGTGTCAGATATATACATCCTG 3'

Position 11: 2 modifications phased 11

5'CTGCAGAAGCTTGGTCCCGGGGCCGCTCAATTGGTCGTAGCAAGCTCTGGATC CGCTTGAT**C**GAACGTACG**C**GCTGTCCCCCGCGTTTTAACCGCCAAGGGGATTACT CCCTAGTCTCCAGGCACGTGTCAGATATATACATCCTG 3'

Each strand was synthesized by ligation of three DNA fragments: left fragment, central fragment (underlined, with modified cytosines in bold red) and right fragment. The complementary counterparts of each of the two strands were synthesized by following the same approach.

The left and right fragments of each strand, as well as the unmodified central fragments, were obtained from Sigma Aldrich. The modified central fragments, containing mdC and hmdC in the specified positions (bold red), were synthesized by solid phase synthesis using the same procedure used for the synthesis of hmdC-modified 8mer and 12mer DNAs for NMR studies, with the following variations: Commercially available 5'-*O*-DMT-dC$^{Ac}$-3'-succinyl-LCAA-CPG (Link Technologies) was used as the solid support. For the synthesis of mdC-modified strands, the 5-Me-dC$^{AC}$-CE phosphoramidite was used (Link Technologies). The coupling time for the 5-Me-dC$^{AC}$-CE phosphoramidite was 15 min. The hmdC-modified and mdC-modified central parts were synthesized in the DMT-OFF mode. In the case of the hmdC-modified strands, the solid supports were treated as in the synthesis of 8mers for NMR studies and purified by 20% denaturing polyacrylamide gel. In the case of mdC-modifed strands, the solid supports were transferred to screw-cap vials and incubated at 55 ºC for 16 h with 1 mL of $NH_3$ solution (30%). After cleavage from the solid support and deprotection, the supernatants were transferred into 2 mL Eppendorf tubes and the supports were rinsed with water (2 x 0.25 mL). The combined solutions were evaporated to dryness using an evaporating centrifuge. The residues that were obtained were purified by 20% denaturing polyacrylamide gel.

To generate the 147 bp 601 Widom fragment, the oligonucleotides for the upper strand (Rec601_For1, 2 and 3) and the ones for the lower strand (Rec601_Rev1, 2 and 3), were phosphorylated using T4 PNK (NEB biolabs) and annealed 2 by 2 in 3 independent reactions by heating 5' at 95ºC and cooling down to RT overnight. The 3 double stranded fragments were then ligated 4hrs at 22ºC using T4 Ligase (NEB biolabs),

| Name | Sequence : (5' to 3') |
| --- | --- |

| | |
|---|---|
| | CTGCAGAAGCTTGGTCCCGGGGCCGCTCAATTGGTCGTAGC |
| Rec601_For1 | AAGCTCTGGATC |
| Rec601_For2 | CGCTTGATCGAACGTACGCGCT |
| | GTCCCCCGCGTTTTAACCGCCAAGGGGATTACTCCCTAGTCT |
| Rec601_For3 | CCAGGCACGTGTCAGATATATACATCCTG |
| | CAGGATGTATATATCTGACACGTGCCTGGAGACTAGGGAGT |
| Rec601_Rev1 | AATCCCCTTGGCGGTTAAAACGCGGG |
| Rec601_Rev2 | GGACAGCGCGTACGTTCGATCA |
| | AGCGGATCCAGAGCTTGCTACGACCAATTGAGCGGCCCCGG |
| Rec601_Rev3 | GACCAAGCTTCTGCAG |

The complete147bp double stranded fragment was purified on 12% polyacrylamide gel and labeled using ($\gamma$–$^{32}$ P)-ATP .

**Supplementary Table S1.** Mass spectrometry analysis of synthesized oligonucleotides*

| Sequence | MW calcd. | MW found |
|---|---|---|
| 5'-dGAAAAAACGGGhmCGAAAAACGG-3' | 6585.0 (+ $Na^+$) | 6582.6 (+ $Na^+$) |
| 5'-dTCCCGTTTTThmCGCCCGTTTTT-3' | 6328.0 (+ $Na^+$) | 6324.7 (+ $Na^+$) |
| 5'-dCGAhmCGTCG-3' | 2433.0 | 2438.3 |
| 5'-dCGCGThmCGACGCG-3' | 3666.3 | 3677.4 |
| CGCTTGAThmCGAAhmCGTACGCGCT | 6750.4 | 6762.6 |
| GGACAGCGCGTAhmCGTThmCGATCA | 6799.4 | 6811.5 |
| CGCTTGAThmCGAACGTACGhmCGCT | 6750.4 | 6762.8 |
| GGACAGhmCGCGTACGTThmCGATCA | 6799.4 | 6811.9 |
| CGCTTGATmCGAAmCGTACGCGCT | 6718.4 | 6729.4 |
| GGACAGCGCGTAmCGTTmCGATCA | 6767.4 | 6779.3 |
| CGCTTGATmCGAACGTACGmCGCT | 6718.4 | 6730.8 |
| GGACAGmCGCGTACGTTmCGATCA | 6767.4 | 6779.0 |

*MALDI-TOF spectra were performed using a Perspective Voyager DETMRP mass spectrometer, equipped with nitrogen laser at 337 nm using a 3 ns pulse. The matrix used contained 2,4,6-

trihydroxyacetophenone (THAP, 10 mg/mL in $CH_3CN$/water 1:1) and ammonium citrate (50 mg/mL in water).

Each double stranded DNA sequence was then incubated with purified histones to allow in vitro nucleosome reconstitution.

Nucleosome reconstitution was performed by the salt dialysis following a similar procedure as described in Perez et al. (3). All DNA and histones to be used in the reactions were freshly quantified immediately prior to use. Briefly, 50 ng of the respective forms of the 601 double stranded fragment (unmethylated control, CpG methylated, and full cytosine methylated) mixed with 2450 ng of carrier DNA were brought to 2M NaCl by adding an equal volume of 4M NaCl. The DNA was further mixed with histones at histone:DNA ratios 1:1 (w/w). The final volume of the reaction was brought to 25 µl using 2M NaCl / 50mM Tris pH 8.0 /1mM EDTA.

Each reconstitution reaction was mixed and transferred to a dialysis chamber (membrane 3,500 MWCO, Pierce). An initial volume of 200 ml of 2M NaCl /50mM Tris pH 8.0 / 1mM EDTA was diluted to 0.2M NaCl with continual addition of 50mM Tris pH 8.0 / 1mM EDTA to a final volume of 2 L using a peristaltic pump set at a flow rate of 40-60ml / hr at 4ºC. The dialyzed reaction was transferred to a microtube and stored at 4ºC.

***Gel mobility shift assays***

Nucleosome reconstitution was analyzed on 6% native polyacrylamide gels, which were pre-electrophoresed for 1hour at 100V at 4ºC in TBE. A 30% sucrose solution was added to the reconstitution reactions as a loading buffer immediately prior to loading the gel. The gels were run at 40 V for 6 hours at 4ºC, dried and exposed to a phosphorimager screen. The band intensities were measured by densitometry using the *PhosphorImager* system (GE Healthcare) (see Figure S).

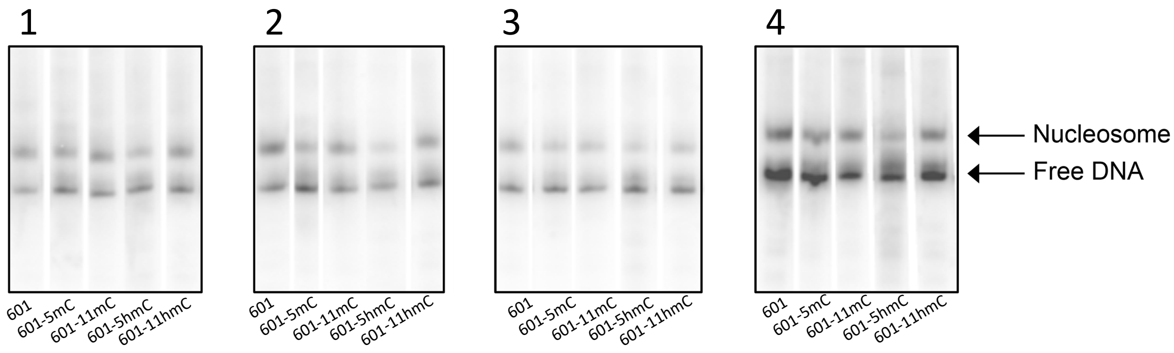


**Supplementary Figure S1**. In vitro nucleosome core particle reconstitution. Results of the four replicas of the gel mobility shift assays of nucleosomes reconstituted in vitro with a 147-bp 601 (with normal cytosines, methylated and hydroxylated respectively). The upper bands (Nucleosome) correspond to histone core-bound DNA, and lower bands correspond to unbound DNA (free DNA). Mk: DNA ladder for size band estimation.

**NMR**

Quantitative distance constraints were obtained from NOESY experiments by using a complete relaxation matrix analysis with the program MARDIGRAS. Error bounds in the interprotonic distances were estimated by carrying out several MARDIGRAS calculations with different initial models, mixing times and correlation times. Standard A- and B-form duplexes were used as initial models, and three correlation times (1.0, 3.0 and 7.0 ns) were employed, assuming an isotropic motion for the molecule. Experimental intensities were recorded at two different mixing times (150 and 250 ms). Final constraints were obtained by averaging the upper and lower distance bounds in all the MARDIGRAS (4) runs. Qualitative limits of 1.8 Å and 5 Å were set in those distances where no quantitative analysis could be carried out, such as overlapping cross-peaks or those with a very weak intensity. In addition to these experimentally derived constraints, Watson-Crick hydrogen bond restraints were used. Target values for distances and angles related to hydrogen bonds were set as described from crystallographic data. No backbone angle constraints were employed. Distance constraints with their corresponding error bounds were incorporated into the AMBER potential energy by defining a flat-well potential term.

**Thermodynamic Integration**

List of all MDB protein-DNA complexes that were subject to thermodynamic integration calculations to establish the differential free energy of binding for methylated and hydroxymethylated DNA.

| MDB | Starting structure | DNA sequence |
|---|---|---|
| MBD1 | PDB ID 1GI4 | GTATC**mC**GGATAC |
| MBD2 | PDB ID 2KY8 | GGAAT**mC**GGCTC |

| | | |
|---|---|---|
| MBD3 | Homology modelling<br>PDB ID 2MB7 (protein) | GGCGCT**mC**GGCGGC |
| MeCP2 | PDB ID 3C2I | ATAGAAGAATTC**mC**GTTCCAG |


#### Supplementary Results

**Supplementary Table S2**. Melting Temperatures (Tm in ºC) for the oligonucleotide CGAC*GTCG, where C* stands for cytosine (C), methylated-cytosine (mC) and hydroxymethylated-cytosine (hmC) respectively, calculated by UV experiments at different oligo concentrations.

| 5’ – CGAC*GTCG - 3’<br>3’ – GCTGC*AGC – 5’ | Tm (ºC) | | |
|---|---|---|---|
| oligonucleotide concentration (μM) | C*=C | C*=mC | C*=hmC |
| 5 | 44.7 | 47.9 | 46.8 |
| 20 | 48.0 | 51.9 | 50.9 |

| | | | |
|---|---|---|---|
| 66 | 51.2 | 55.7 | 54.1 |

**Supplementary Table S3**. Variation in thermodynamic parameters, enthalphy (ΔH in Kcal/mol), entropy (ΔS in cal/mol*K) and free energy ΔG (Kcal/mol), at 25 ºC calculated using Van't Hoff equation (see Methods) from UV data for he oligomer in Table 1 with cytosine, methyl-cytosine and hydroxymethycytosine respectively.

| | **C** | **mC** | **hmC** |
|---|---|---|---|
| ΔH (Kcal/mol) | -82.4 | -70.2 | -73.1 |
| ΔS (cal/mol*K) | -232.4 | -191.8 | -201.5 |
| ΔG (at 25 ºC) (Kcal/mol) | -13.2 | -13.1 | -13.1 |

**Supplementary Table S4**: Assignment of the proton resonances of the hMC duplex $(CGAhCGTCG)_2$, where hC = 5-hydroxymethylcytosine. Buffer conditions: 25 mM sodium phosphate, 100 mM NaCl, T= 5 ºC, pH 7.

| | H1' | H2'/H2" | H3' | H4' | H5'/H5" | H5/Met | H6/H8 | H1/H3 |
|---|---|---|---|---|---|---|---|---|
| **C1** | 5.66 | 1.92/2.38 | 4.71 | 4.06 | 3.73 | 5.87 | 7.63 | 7.12/8.30 |
| **G2** | 5.52 | 2.78 | 5.04 | 4.35 | 4.12/3.98 | -- | 7.99 | 12.99 |
| **A3** | 6.29 | 2.66/2.98 | 5.04 | 4.51 | 4.31/4.20 | 7.91 | 8.26 | -- |
| **hC4** | 5.51 | 2.12/2.40 | 4.84 | 4.19 | 4.30 | 3.99 | 7.28 | 6.49/8.54 |
| **G5** | 5.97 | 2.61/2.81 | 4.95 | 4.40 | 4.24/4.16 | -- | 7.86 | 12.71 |
| **T6** | 6.07 | 2.13/2.49 | 4.91 | 4.26 | 4.15 | 1.35 | 7.34 | 13.87 |
| **C7** | 5.67 | 2.05/2.39 | 4.87 | 4.15 | 4.09 | 5.68 | 7.51 | 7.18/8.68 |
| **G8** | 6.15 | 2.36/2.65 | 4.71 | 4.21 | 4.10 | -- | 7.97 | 13.23 |

**Supplementary Table S5**: Assignment of the proton resonances of the 5MC duplex $(CGAmCGTCG)_2$, where mC = 5-methylcytosine. Buffer conditions: 25 mM sodium phosphate, 100 mM NaCl, T= 5 ºC, pH 7.

| | H1' | H2'/H2" | H3' | H4' | H5'/H5" | H5/Met | H6/H8 | H1/H3 |
|---|---|---|---|---|---|---|---|---|
| **C1** | 5.71 | 1.95/2.40 | 4.72 | 4.07 | 3.73 | 5.92 | 7.66 | 7.18/8.30 |
| **G2** | 5.52 | 2.78 | 5.02 | 4.34 | 4.10 | -- | 7.99 | 13.00 |
| **A3** | 6.31 | 2.70/3.02 | 5.04 | 4.51 | 4.30/4.18 | 7.92 | 8.31 | -- |
| **mC4** | 5.56 | 2.09/2.38 | 4.84 | 4.17 | 4.30 | 1.60 | 7.08 | 6.23/8.46 |
| **G5** | 5.95 | 2.60/2.78 | 4.92 | 4.38 | 4.25/4.14 | -- | 7.78 | 12.80 |
| **T6** | 6.09 | 2.13/2.50 | 4.90 | 4.26 | 4.15 | 1.34 | 7.34 | 13.89 |
| **C7** | 5.70 | 2.04/2.39 | 4.87 | 4.12 | -- | 5.69 | 7.52 | 7.19/8.70 |
| **G8** | 6.18 | 2.36/2.64 | 4.71 | 4.21 | 4.08 | -- | 7.99 | 13.22 |

**Supplementary Table S6**: Assignment of the proton resonances of the control duplex $(CGACGTCG)_2$. Buffer conditions: 25 mM sodium phosphate, 100 mM NaCl, T= 5 ºC, pH 7.

| | H1′ | H2′/H2″ | H3′ | H4′ | H5′/H5″ | H5/Met | H6/H8 | H1/H3 |
|---|---|---|---|---|---|---|---|---|
| **C1** | 5.71 | 1.94/2.40 | 4.72 | 4.06 | 3.73 | 5.93 | 7.65 | 7.20/8.33 |
| **G2** | 5.48 | 2.78 | 5.03 | 4.33 | 4.10 | -- | 8.01 | 13.03 |
| **A3** | 6.29 | 2.74/2.96 | 5.08 | 4.52 | 4.28/4.19 | 7.91 | 8.26 | -- |
| **C4** | 5.58 | 2.05/2.37 | 4.85 | 4.18 | 4.28 | 5.25 | 7.25 | 6.64/8.21 |
| **G5** | 5.97 | 2.63/2.80 | 4.96 | 4.39 | 4.25/4.13 | -- | 7.87 | 12.80 |
| **T6** | 6.06 | 2.10/2.48 | 4.88 | 4.25 | 4.13 | 1.41 | 7.31 | 13.91 |
| **C7** | 5.71 | 2.06/2.39 | 4.86 | 4.12 | -- | 5.71 | 7.53 | 7.20/8.71 |
| **G8** | 6.19 | 2.37/2.65 | 4.71 | 4.21 | 4.10 | -- | 7.99 | 13.23 |

**Supplementary Table S7**: Assignment of the proton resonances of the hMC duplex d(CGCGAhCGTCGCG)$_2$. Buffer conditions: 25 mM sodium phosphate, 100 mM NaCl, T= 5 ºC, pH 7.

| | H1′ | H2′/H2″ | H3′ | H4′ | H5′/H5″ | H5/Met/H2 | H6/H8 | H1/H3/H41/2 |
|---|---|---|---|---|---|---|---|---|
| **C1** | 5.77 | 1.99/2.43 | 4.72 | 4.07 | 3.73 | 5.92 | 7.63 | 7.16/8.20 |
| **G2** | 5.92 | 2.67/2.76 | 4.99 | 4.36 | 4.1/3.99 | -- | 7.97 | 13.11 |
| **C3** | 5.72 | 2.10/2.44 | 4.86 | 4.21 | 4.11/4.20 | 5.40 | 7.35 | 6.55/8.37 |
| **G4** | 6.00 | 2.66/2.79 | 4.99 | 4.39 | 4.08/4.21 | -- | 7.92 | 12.9 |
| **T5** | 6.00 | 1.92/2.46 | 4.85 | 4.22 | 4.10 | 1.42 | 7.18 | 13.83 |
| **hC6** | 5.48 | 2.01/2.34 | 4.84 | -- | -- | 4.09/4.22 | 7.45 | 6.61/8.82 |
| **G7** | 5.51 | 2.78/2.69 | 5.01 | 4.34 | -- | -- | 7.92 | 12.7 |
| **A8** | 6.19 | 2.65/2.88 | 5.03 | 4.45 | 4.14/4.20 | 7.81 | 8.18 | -- |
| **C9** | 5.56 | 1.89/2.29 | 4.84 | -- | -- | 5.22 | 7.18 | 6.59/8.18 |
| **G10** | 5.84 | 2.59/2.69 | 4.96 | 4.34 | 4.00/4.10 | -- | 7.84 | 12.96 |
| **C11** | 5.75 | 1.90/2.33 | 4.96 | 4.16 | 4.10 | 5.42 | 7.31 | 6.70/8.49 |
| **G12** | 6.17 | 2.60/2.35 | 4.68 | 4.07 | 4.18 | -- | 7.93 | 13.11 |

**Supplementary Table S8**: Assignment of the proton resonances of the control duplex d(CGCGACGTCGCG)$_2$. Buffer conditions: 25 mM sodium phosphate, 100 mM NaCl, T= 5 ºC, pH 7.

| | H1′ | H2′/H2″ | H3′ | H4′ | H5′/H5″ | H5/Met/H2 | H6/H8 | H1/H3/H41/2 |
|---|---|---|---|---|---|---|---|---|
| **C1** | 5.75 | 2.04/2.45 | 4.73 | 4.08 | 3.73 | 5.92 | 7.67 | 7.18/8.21 |
| **T2** | 5.92 | 2.69/2.76 | 4.99 | 4.37 | 4.00/4.10 | -- | 8.00 | 13.12 |
| **A3** | 5.71 | 2.13/2.45 | 4.87 | 4.22 | 4.16 | 5.40 | 7.37 | 6.57/8.39 |
| **C4** | 6.00 | 2.65/2.82 | 4.99 | 4.40 | 4.09/4.23 | -- | 7.96 | 12.92 |
| **G5** | 5.99 | 2.09/2.47 | 4.85 | 4.22 | 4.13 | 1.41 | 7.24 | 13.83 |
| **C6** | 5.54 | 1.98/2.36 | 4.83 | -- | 4.07 | 5.59 | 7.43 | 6.93/8.53 |
| **G7** | 5.53 | 2.79/2.70 | 5.00 | 4.33 | 4.01/4.09 | -- | 7.92 | 12.76 |
| **C8** | 6.17 | 2.64/2.88 | 5.01 | 4.45 | 4.10/4.20 | 7.77 | 8.18 | -- |
| **G9** | 5.53 | 1.93/2.29 | 4.81 | 4.22 | 4.11 | 5.18 | 7.20 | 6.60/8.20 |
| **T10** | 5.85 | 2.59/2.69 | 4.96 | 4.35 | 4/4.10 | -- | 7.87 | 12.96 |
| **A11** | 5.71 | 1.93/2.34 | 4.83 | 4.16 | 4.10 | 5.43 | 7.35 | 6.72/8.49 |
| **G12** | 6.17 | 2.64/2.35 | 4.69 | 4.07 | 4.19 | -- | 7.97 | 13.12 |

**Supplementary Table S9**: NMR restraints and structural calculation statistics.

| | hMC Duplex | 5MC Duplex | hMC 12-mer | Control |
|---|---|---|---|---|
| **Experimental distance constraints** | | | | |
| Total number | 246 | 202 | 302 | 326 |
| Intra-residual | 134 | 124 | 124 | 148 |
| Sequential | 94 | 64 | 132 | 152 |
| Inter-strand | 18 | 14 | 46 | 26 |
| **RMSD** (Å) | | | | |
| Backbone atoms | 1.0 ± 0.2 Å | 0.9 ± 0.2 Å | 1.0 ± 0.3 Å | 0.8± 0.2 Å |
| Base heavy atoms | 0.6 ± 0.1 Å | 0.6 ± 0.2 Å | 0.6 ± 0.2 Å | 0.5 ± 0.2 Å |
| All heavy atoms | 0.9 ± 0.2 Å | 0.9 ± 0.2 Å | 0.9 ± 0.2 Å | 0.7 ± 0.2 Å |
| **Residual violations** | Average (range) | | | |
| Sum of violations (Å) | 55.0 (53.8 - 59.2) | 38.6 (38.1- 40.0) | 15.8 (14.0-18.0) | 4.0 (3.5-4.9) |
| Max. violation (Å) | 0.3 (0.4 - 0.2) | 0.3 (0.4 - 0.2) | 0.44 (0.30-0.51) | 0.32 (0.25-0.39) |
| NOE energy (kcal/mol) | 55.8 (53.8 - 59.2) | 65.6 (63.3 - 67.0) | 69.8 (66.5-72.1) | 15.5 (10.1-20.2) |

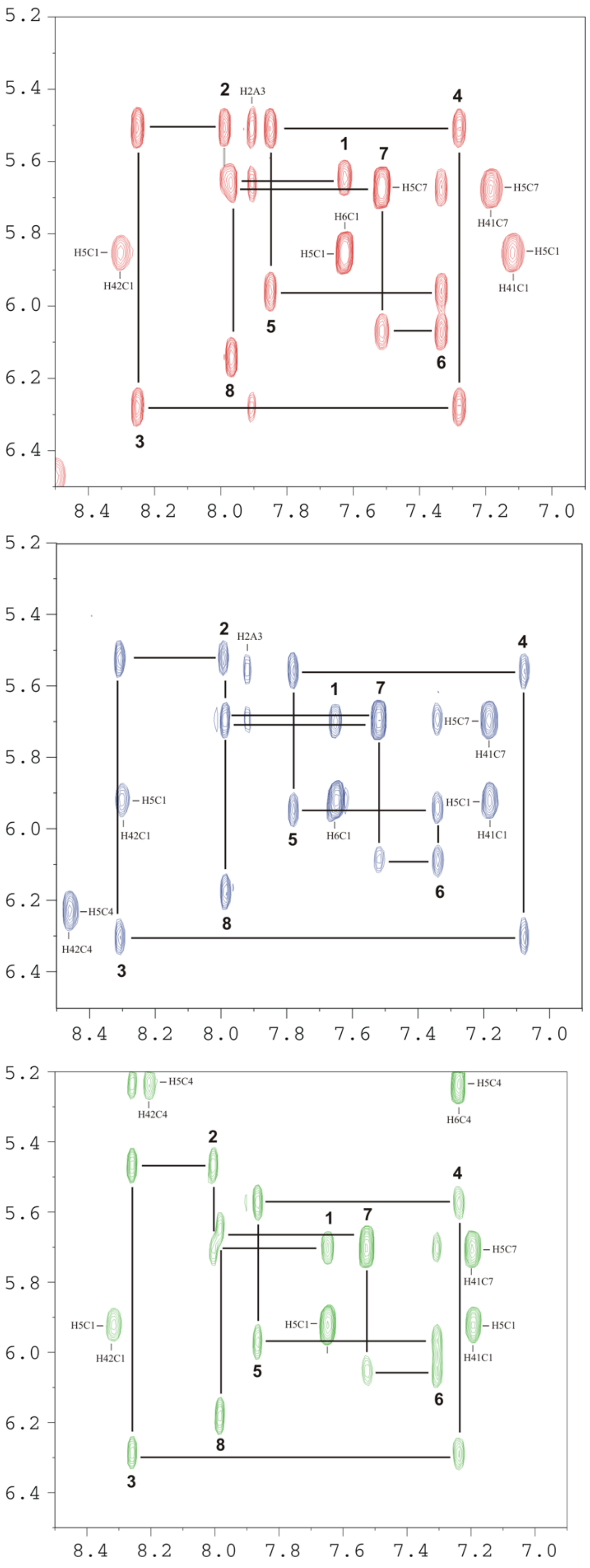


**Supplementary Figure S2**: Regions of NOESY spectra (150 ms mixing time) of hMC $(CGA*CGTCG)_2$ duplexes, *C= hMC (top), 5MC duplex (middle) and C (control) duplex (bottom). H1'-base assignment pathways are indicated. Buffer conditions: 100 mM NaCl, 25 mM sodium phosphate, T=5ºC, pH 7.

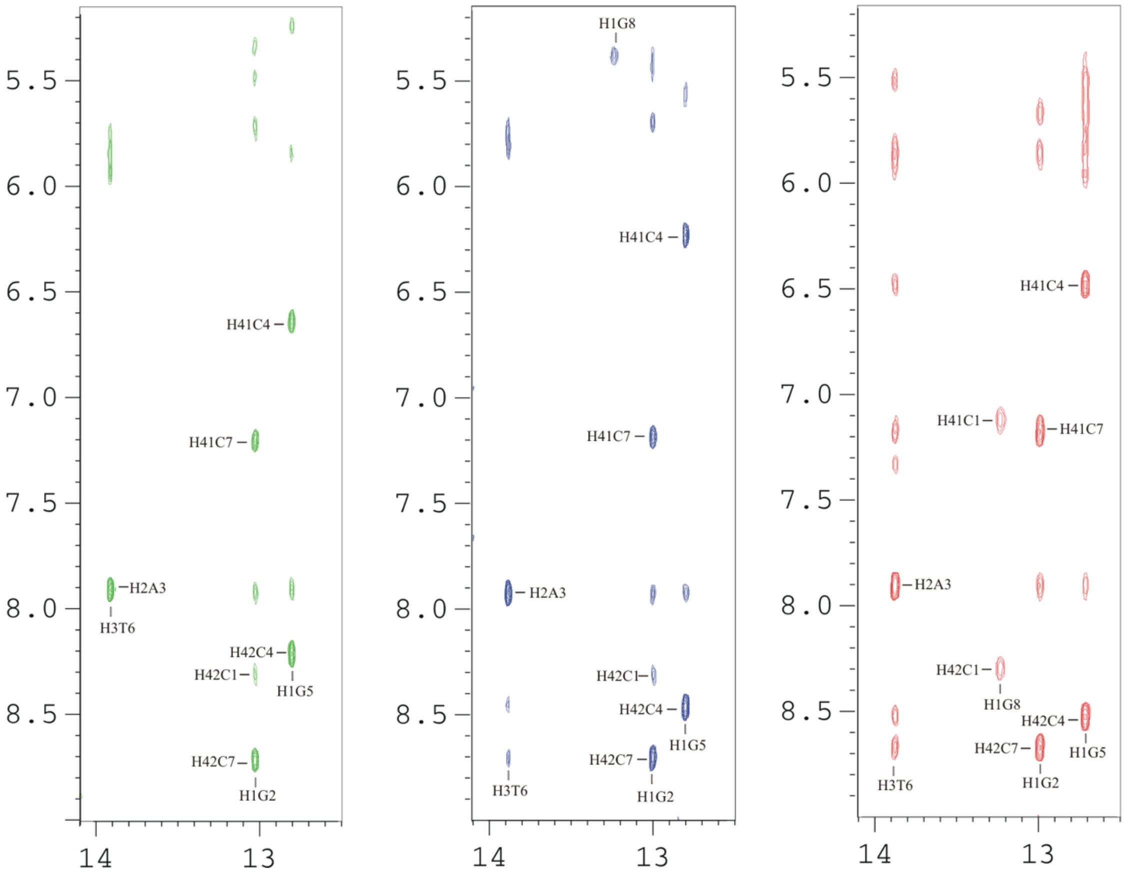


**Supplementary Figure S3**: Imino region of the NOESY spectra in $H_2O$ (τm=150 ms) of $(CGA*CGTCG)_2$ duplexes, *C=hMC (right), 5MC (middle), and C (control) duplex (left). Buffer conditions: 100 mM NaCl , 25mM sodium phosphate, T=5 ºC, pH 7.

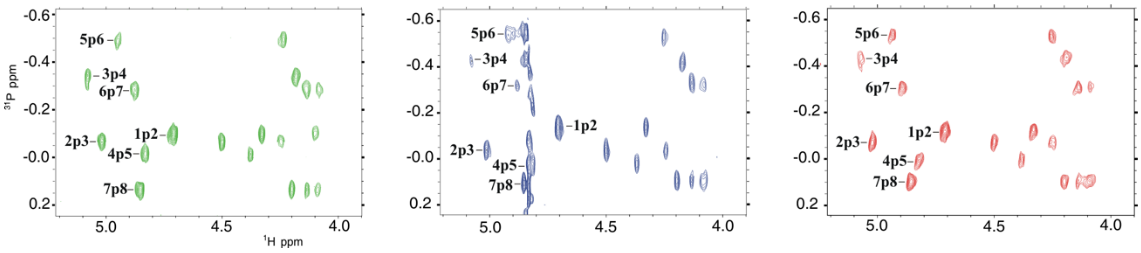

**Supplementary Figure S4**: $^{1}H$-$^{31}P$ correlation spectra for hMC (right), 5MC (middle), and control duplex (left). Buffer conditions were 100 mM NaCl , 25mM sodium phosphate, T=5 ºC, pH 7.

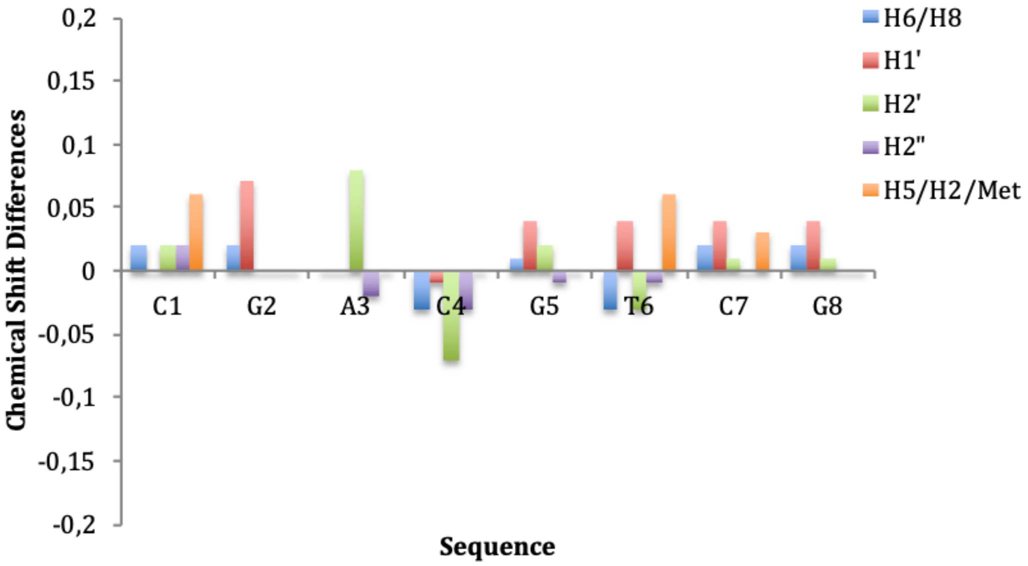


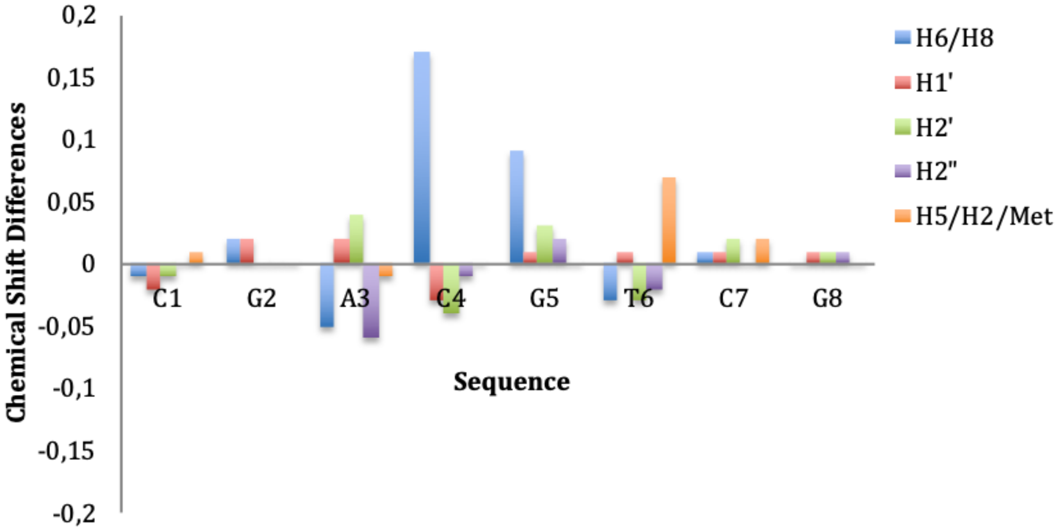


**Supplementary Figure S5**: Chemical shifts differences for non-exchangeable protons between hMC (top) and 5MC (bottom) with respect to the control duplex.

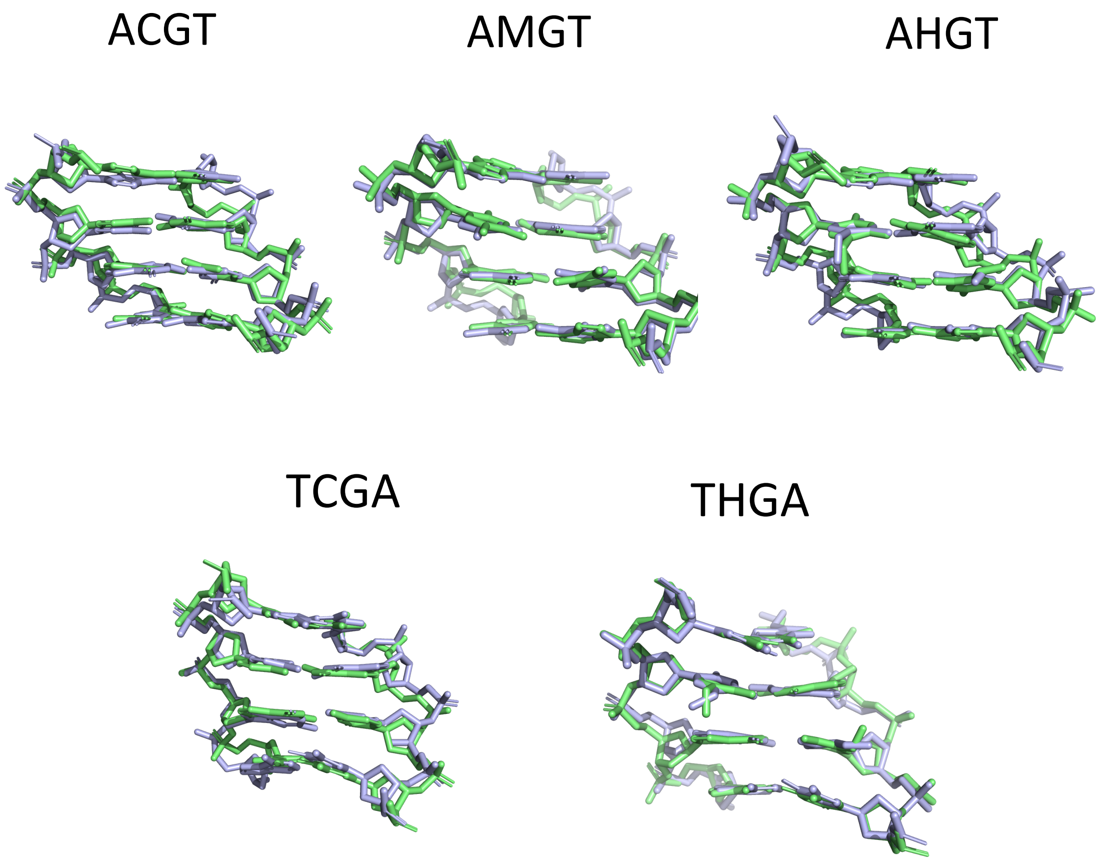


**Supplementary Figure S6.** Overlap of the central tetramer of the average NMR structures (light blue) with the average structure from MD (green), for the tetramers ACGT, AMGT, AHGT, TCGA and THGA. RMDs calculated between the structures for heavy atoms 0.95, 0.69, 0.98, 1.1, 0.76 Å respectively.

**Supplementary Table S10.** Average parameters (in Å and Degrees) averaged over the last 200 ns for the central step (d(C*pG)·d(C*pG)) in the different tetrameric environments between the different forms of cytosine, HydroxyMethylC, MethylC, Cytosine.

| | A-CG-A | A-CG-C | A-CG-G | A-CG-T | C-CG-A | C-CG-C | C-CG-G | G-CG-A | G-CG-C | T-CG-A | AVG | SD_TOT |
|---|---|---|---|---|---|---|---|---|---|---|---|---|
| SHIFT | 0.63±0.97 | -0.19±0.96 | 0.19±1.08 | -0.13±0.90 | 0.34±0.99 | 0.16±0.95 | 0.06±0.95 | 0.56±0.89 | 0.08±0.98 | -0.11±0.99 | 0.16 | 0.28 |
| SLIDE | 0.23±0.52 | -0.03±0.51 | 0.17±0.54 | -0.05±0.50 | 0.23±0.55 | 0.06±0.55 | 0.06±0.58 | 0.18±0.54 | 0.01±0.49 | 0.14±0.55 | 0.10 | 0.11 |
| RISE | 3.28±0.34 | 3.39±0.36 | 3.29±0.33 | 3.59±0.34 | 3.07±0.33 | 3.09±0.35 | 3.01±0.33 | 3.12±3.01 | 3.20±0.34 | 3.08±0.35 | 3.21 | 0.18 |
| TILT | 3.70±5.69 | -0.59±5.96 | 0.60±6.22 | -0.50±5.92 | 2.50±5.47 | 0.70±5.44 | -0.10±5.39 | 3.00±5.17 | 0.17±5.69 | -0.60±5.55 | 0.89 | 1.60 |
| ROLL | 6.90±6.53 | 6.63±6.59 | 5.70±6.49 | 7.20±6.53 | 6.65±6.20 | 6.20±6.17 | 5.50±6.19 | 7.00±6.38 | 6.31±6.36 | 8.30±6.19 | 6.64 | 0.81 |
| TWIST | 35.30±6.61 | 37.32±6.34 | 36.64±6.28 | 41.40±5.20 | 30.90±7.35 | 31.40±7.40 | 29.80±7.52 | 31.50±7.16 | 32.90±7.22 | 29.40±7.84 | 33.66 | 3.89 |

| | A-Mg-A | A-Mg-C | A-Mg-G | A-Mg-T | C-Mg-A | C-Mg-C | C-Mg-G | G-Mg-A | G-Mg-C | T-Mg-A | AVG | SD_TOT |
|---|---|---|---|---|---|---|---|---|---|---|---|---|
| SHIFT | 0.42±0.85 | -0.06±0.88 | 0.20±0.86 | -0.06±0.89 | 0.16±0.75 | 0.10±0.75 | -0.10±0.71 | 0.20±0.75 | 0.02±0.79 | 0.02±0.68 | 0.11 | 0.16 |
| SLIDE | 0.07±0.51 | -0.09±0.49 | -0.01±0.51 | -0.22±0.52 | 0.03±0.49 | -0.02±0.45 | -0.01±0.45 | 0.01±0.45 | -0.07±0.44 | 0.00±0.44 | -0.01 | 0.05 |
| RISE | 3.25±0.36 | 3.32±0.36 | 3.18±0.36 | 3.59±0.37 | 2.98±0.32 | 3.01±0.31 | 2.95±0.30 | 3.04±0.30 | 3.10±0.32 | 2.97±0.31 | 3.09 | 0.13 |
| TILT | 2.47±5.16 | 0.26±5.23 | 0.69±5.06 | -0.11±5.79 | 1.64±0.45 | -0.22±4.54 | -0.02±4.51 | 0.94±4.46 | -0.05±4.54 | 0.08±4.41 | 0.65 | 0.90 |
| ROLL | 12.27±6.78 | 11.18±6.72 | 11.04±6.42 | 11.25±6.53 | 11.58±5.9 | 11.41±5.92 | 11.18±5.84 | 12.54±5.86 | 11.90±6.25 | 13.12±5.66 | 11.80 | 0.71 |
| TWIST | 30.56±7.31 | 32.72±6.69 | 30.67±7.15 | 37.76±6.02 | 25.43±7.07 | 27.01±6.55 | 26.43±6.43 | 25.64±6.54 | 27.53±6.58 | 23.07±6.45 | 27.67 | 3.07 |

| | A-HJ-A | A-HJ-C | A-HJ-G | A-HJ-T | C-HJ-A | C-HJ-C | C-HJ-G | G-HJ-A | G-HJ-C | T-HJ-A | AVG | SD_TOT |
|---|---|---|---|---|---|---|---|---|---|---|---|---|

| | | | | | | | | | | | | |
|---|---|---|---|---|---|---|---|---|---|---|---|---|
| SHIFT | 0.17±0.58 | 0.09±0.57 | 0.13±0.55 | -0.05±0.85 | 0.01±0.51 | 0.02±0.52 | -0.03±0.5 | 0.10±0.51 | 0.03±0.55 | -0.05±0.69 | 0.06 | 0.07 |
| SLIDE | 0.00±0.45 | -0.07±0.46 | -0.04±0.44 | -0.23±0.50 | -0.02±0.41 | -0.05±0.42 | -0.05±0.39 | 0.02±0.43 | -0.05±0.44 | -0.00±3.02 | -0.03 | 0.03 |
| RISE | 3.06±0.31 | 3.16±0.31 | 3.02±0.29 | 3.59±0.36 | 2.87±0.28 | 2.92±0.29 | 2.81±0.28 | 2.94±0.28 | 3.05±0.30 | 3.03±0.32 | 2.99 | 0.11 |
| TILT | 1.30±4.11 | 0.30±4.00 | 0.10±3.91 | -0.56±6.00 | 1.00±3.95 | -0.15±3.84 | -0.10±3.90 | 0.60±3.90 | -0.05±3.85 | 0.08±4.53 | 0.35 | 0.53 |
| ROLL | 13.70 ± 5.61 | 13.20 ± 5.72 | 12.50 ± 5.39 | 11.12 ± 6.45 | 12.66 ± 5.35 | 12.44 ± 5.38 | 11.13 ± 5.31 | 14.00 ± 5.31 | 13.10 ± 5.41 | 13.56 ± 5.93 | 12.91 | 0.89 |
| TWIST | 27.20 ± 6.33 | 30.80 ± 6.16 | 28.80 ± 6.09 | 38.90 ± 5.8 | 23.50 ± 5.77 | 25.80 ± 6.09 | 24.70 ± 6.09 | 24.30 ± 5.63 | 27.50 ± 6.54 | 24.48 ± 6.85 | 26.34 | 2.41 |

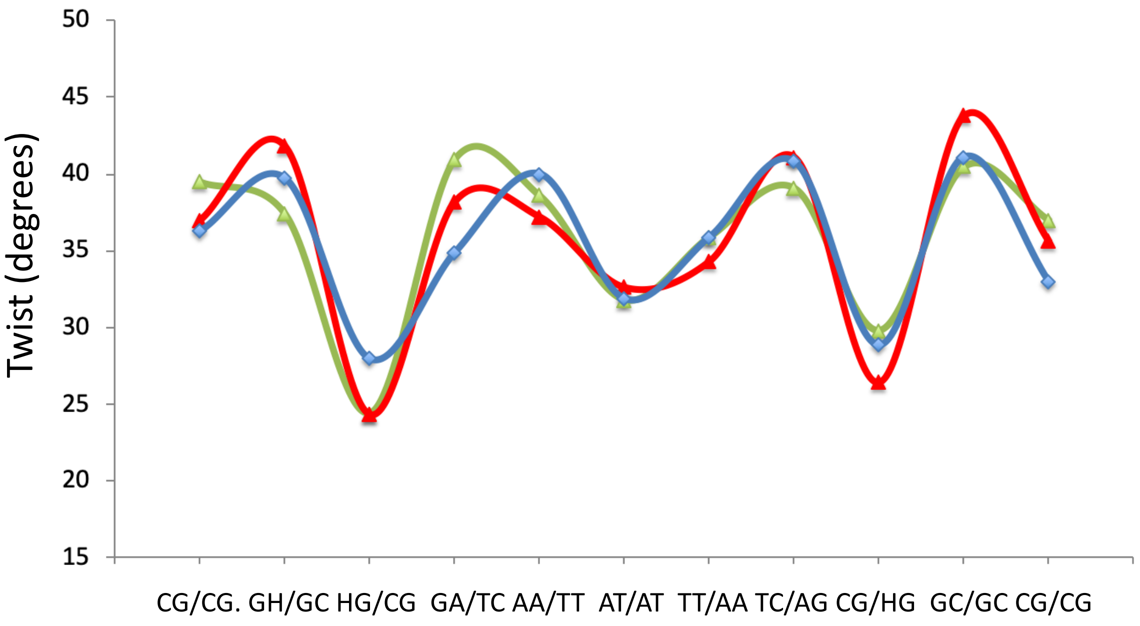


**Supplementary Figure S7.** Twist profile for the hemi-hydroxymethylated sequence in the X-ray crystal structures 4GLH (red line), 4HLI (green line) and 4GLC (blue line). HG/CG steps are characterised by low-twist state.

**Supplementary Table S11.** Diagonal stiffness constants for translational movements in kcal/mol ang$^2$ for the central C*pG step (C*=C, mC and hmC) in the different tetrameric environments.

| | $K^b_{shift\text{-}shift}$ | | | $K^b_{slide\text{-}slide}$ | | | $K^b_{rise\text{-}rise}$ | | |
|---|---|---|---|---|---|---|---|---|---|
| | CpG | mCpG | hmCpG | CpG | mCpG | hmCpG | CpG | mCpG | hmCpG |
| ACGA | 1.28 | 1.29 | 1.96 | 2.91 | 2.88 | 3.31 | 7.91 | 7.79 | 9.09 |
| ACGC | 1.34 | 1.09 | 1.97 | 2.77 | 2.80 | 3.14 | 7.19 | 7.07 | 8.59 |
| ACGG | 1.16 | 1.11 | 2.04 | 2.79 | 2.74 | 3.42 | 7.87 | 7.67 | 9.63 |
| ACGT | 1.57 | 1.31 | 1.50 | 3.14 | 2.72 | 2.85 | 7.28 | 6.69 | 6.65 |
| CCGA | 1.01 | 1.34 | 2.30 | 2.94 | 3.21 | 3.95 | 8.80 | 9.35 | 11.21 |
| CCGC | 1.13 | 1.20 | 2.17 | 2.82 | 3.51 | 3.92 | 8.12 | 9.31 | 10.67 |
| CCGG | 1.03 | 1.27 | 2.40 | 3.02 | 3.32 | 4.29 | 8.41 | 9.46 | 11.54 |
| GCGA | 1.32 | 1.33 | 2.29 | 3.14 | 3.58 | 3.71 | 8.64 | 9.30 | 10.93 |
| GCGC | 1.17 | 1.07 | 1.96 | 3.20 | 3.56 | 3.66 | 8.06 | 8.48 | 9.47 |
| TCGA | 1.01 | 1.42 | 1.38 | 2.91 | 3.51 | 2.98 | 8.63 | 9.47 | 8.99 |
| poli(CG) | 0.022 | 0.024 | 0.027 | 0.018 | 0.015 | 0.018 | 0.033 | 0.030 | 0.030 |

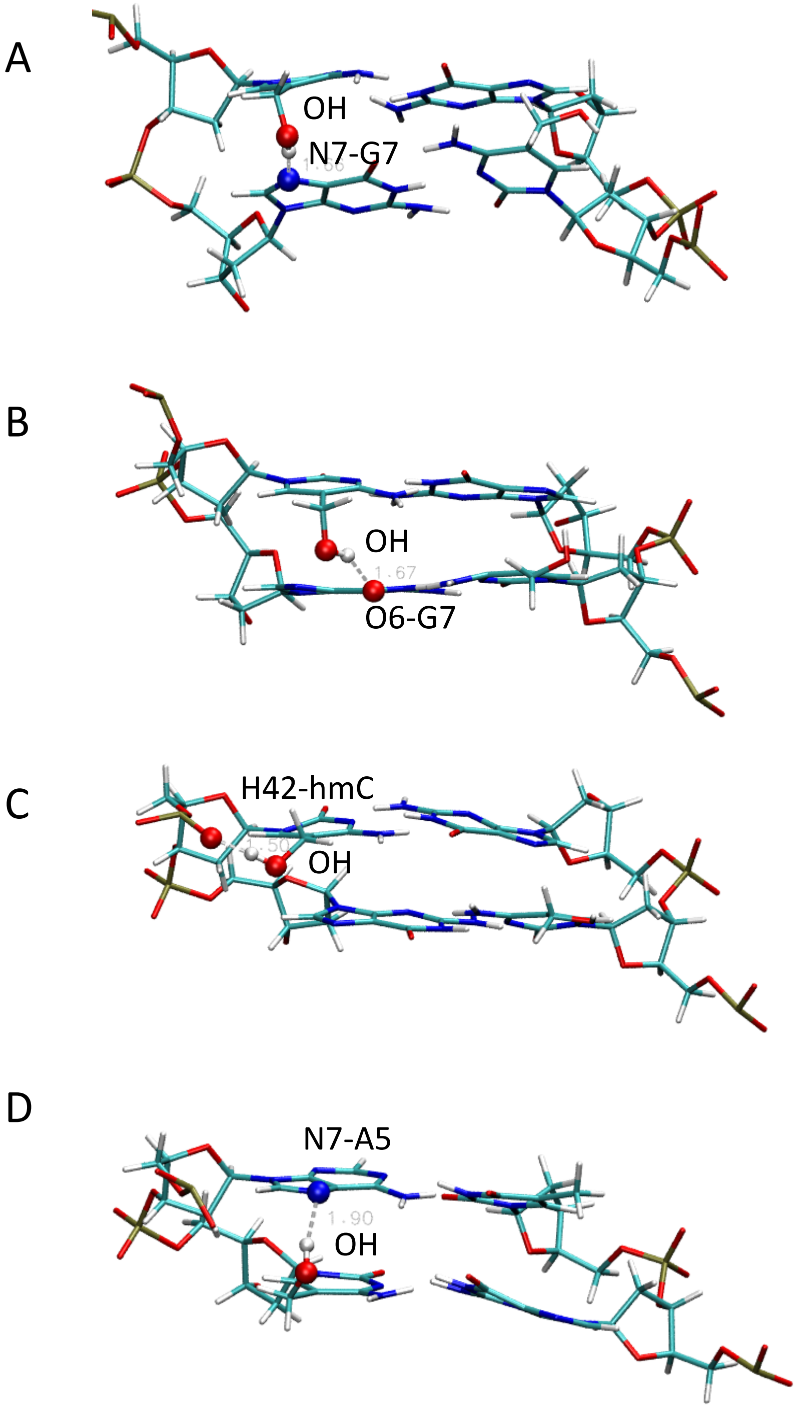


**Supplementary Figure S8.** Hydrogen bonds detected along the MD simulations among hydroxy group of hmC (OH, hmC position 6) and the flanking bases (guanine 7, adenine 5). We detected the formation of hydrogen bonds between the hydrogen of the hydroxyl group and the neighbouring guanine (G7); in particular between the hydrogen of the hydroxyl group and the nitrogen 7 (panel A) or/and the oxygen 6 (panel B) of the guanine. We also detected a HB between the hydroxyl hydrogen and the oxygen of the backbone (panel C). It is worth noticing that these hydrogen bonds are formed maximum the 0.039 fraction of the simulation time and they are not stable. Nevertheless, the interaction between hmC and G7 indicates that the base pair can assume a more distorted conformation, that reflects into an opening of the minor groove, high roll. In the AC*GT tetramer (panel D) the hydroxyl group interacts also with the adenine (A6), and not only with the following guanine. This behaviour translates into the stiffer and higher twist of the tetramers XCGY where X is a purine and Y a pyrimidine base.

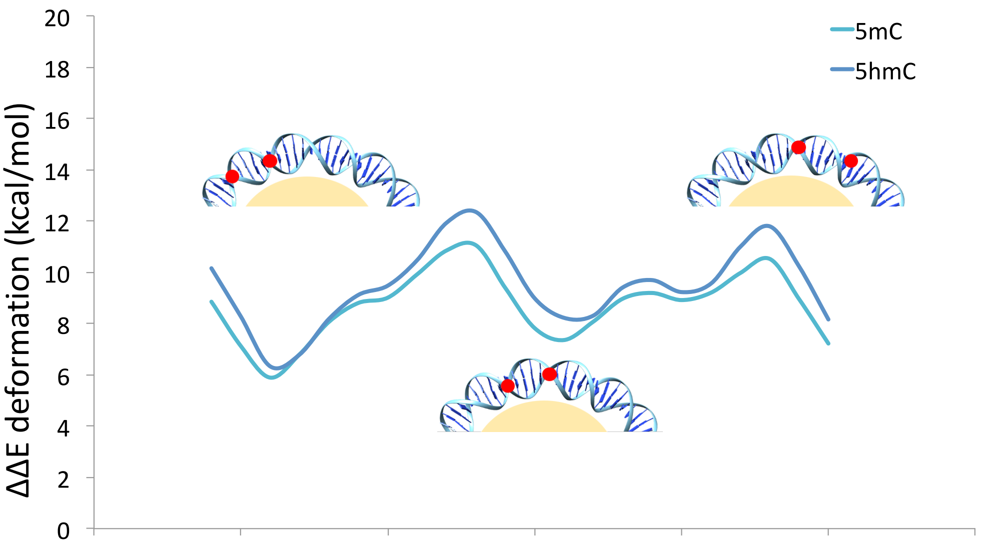


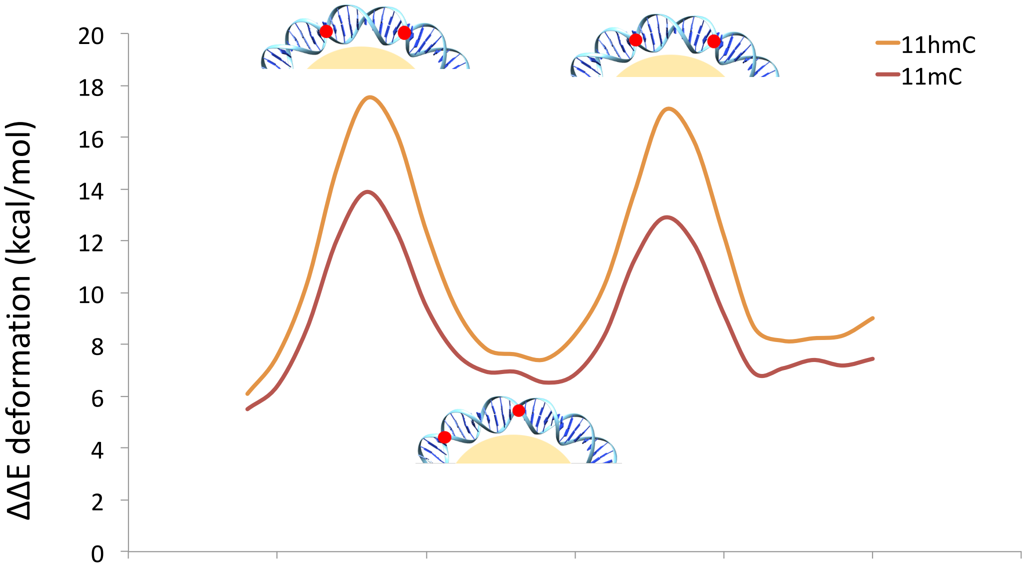

**Supplementary Figure S9.** Periodicity of the variation in deformation energy given by the positioning of the epigenetic modifications respect to the histones in the nucleosome. In the top panel, in blue (light blue mC and dark blue hmC) the variation in energy when the modifications are 5 base pairs apart. IIn the bottom panel, energy variations (light red mC and dark red hmC) when the modifications are 11 base pairs apart. In both plots a schematic image of the DNA modifications (red dots) when they are positioned 5bps or 11 bps apart, in the minor or major grooves facing the histones (in yellow).

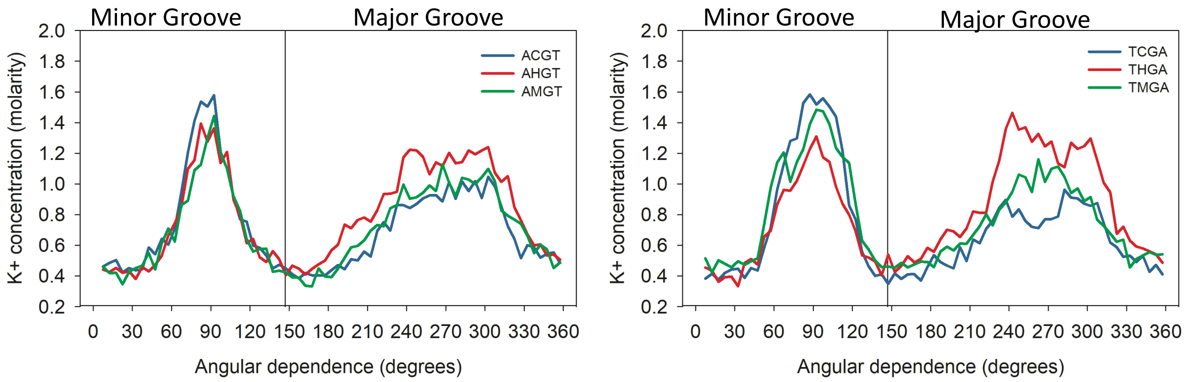


**Supplementary Figure S10**. Cation K+ concentration (molarity) along the minor and major grooves averaged over the last 200 ns of the trajectories. K+ molarity distribution as a function of the angular dependence (in degrees) for the tetramers AC*GT and TC*GA where C*=C,mC,hmC ( in blue, green and red respectively).

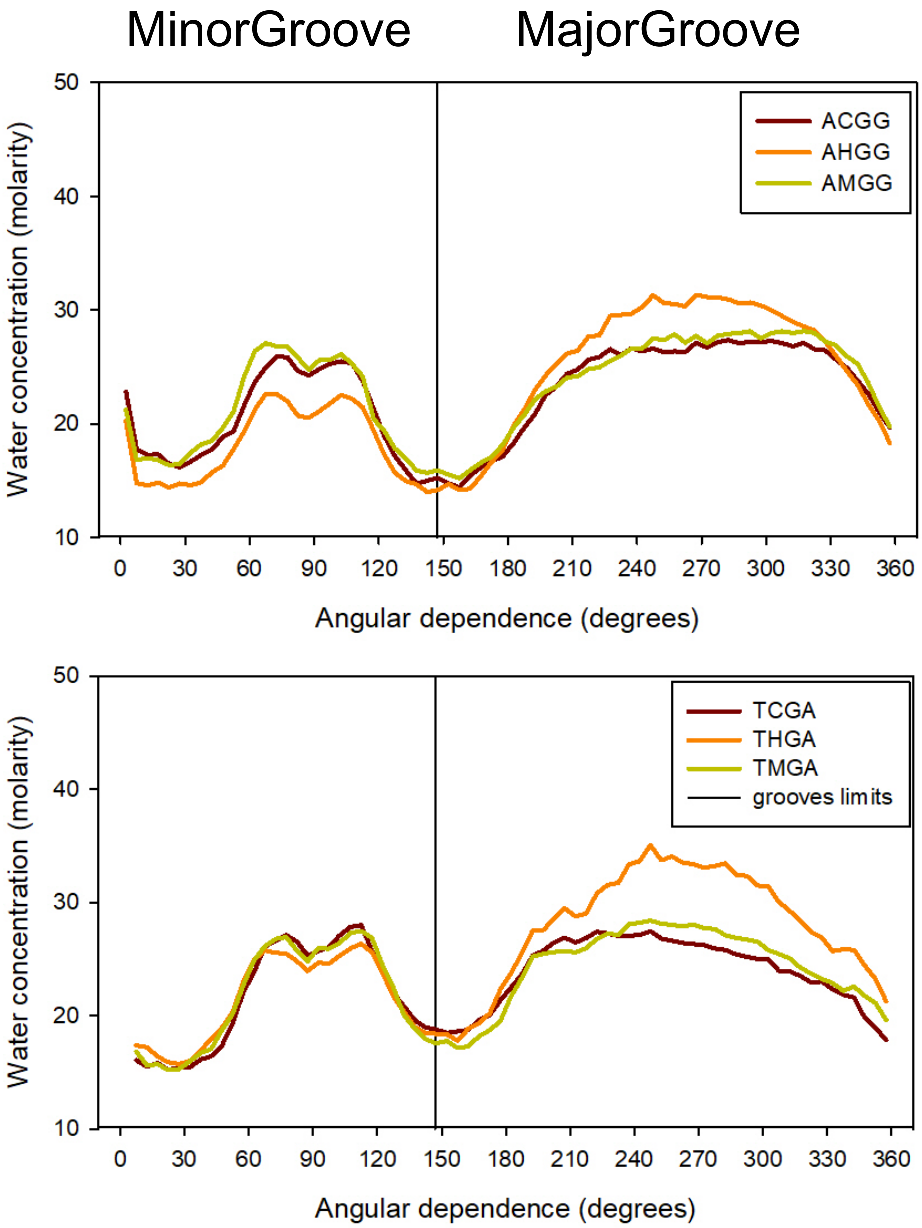


**Supplementary Figure S11.** Occupancy maps of water molecules in the major and minor groove for the unmodified CpG, mCpG and hmCpG respectively in the two tetramer AC*GG and TC*GA.
